# A photoprotection dial maps holistic light-stress response in diatoms

**DOI:** 10.64898/2025.12.11.693581

**Authors:** D. Croteau, M. Jaubert, T. Quemar, A. Falciatore, A. Maes, B. Bailleul

## Abstract

Cellular responses to light stress operate on two timescales: short-term responses, driven by dynamic photosynthetic mechanisms such as non-photochemical quenching (NPQ), and long-term responses, characterized by various physiological and metabolic shifts driven by changes in gene expression. Understanding the coordination within and between these layers represents a major challenge in photosynthesis research. We achieved this in the diatom *Phaeodactylum tricornutum* using an adjustable “photoprotection dial” across 10 mutant strains expressing different constitutive levels of the Lhcx1 protein, which determine their latent NPQ capacity. Crucially, a near-identical initial state is preserved among strains, which allows a precise mapping of the causal chain of events initiated by NPQ induction under high light. This unlocks exceptional leverage to examine holistic interactions between functional photosynthesis and gene expression within their natural regulatory architecture. We observe that increasing NPQ maintains the PSII primary acceptor Q_A_ in a more oxidized state with little effect on linear electron flow. This oxidation limits PSII photodamage and promotes cyclic electron flow around PSI, which in turn is consistent with enhanced ATP production supporting PSII repair. Moreover, transcriptomic analysis across high-light exposure time suggests that the expression of nearly half of the genome is modulated via NPQ-driven effects. Gene expression profile clustering reveals distinct and coherent transcriptomic regulatory networks. The short-term response is characterized by a strong downregulation of light-harvesting-related genes — independently of NPQ — and an upregulation of oxidative stress response genes that scales inversely with NPQ magnitude. In the long term, most of the transcriptome is deeply impacted by high-light stress in a widely NPQ-dependent manner, highlighting the critical role of this process in shaping photophysiology, which in turn influences gene expression. By engineering a native functional feedback loop into an experimental dial, our approach establishes a generalizable framework for studying the complex interplay between physiology and system-wide biological regulation.

## Introduction

Photosynthetic organisms continuously adjust to fluctuating environmental conditions and energy demands, thanks to an integrated network of functional feedback loops. These mechanisms are tightly coupled to photosynthesis and operate within the plastid to modulate the activity of complexes in response to changes in redox poise or proton gradient buildup. For instance, under high-light stress, a decrease in luminal pH activates non-photochemical quenching (NPQ) (Goss and Lepetit, 2015), which in turn reduces photosystem (PS) II efficiency. Such rapid responses represent what (Bich et al., 2016) termed "dynamic stability", an immediate frontline response that confers robustness to photosynthesis during perturbations. Simultaneously, photosynthesis generates signals that influence nuclear gene expression through retrograde signalling (Chan et al., 2016; Escoubas et al., 1995; Jung et al., 2013; Lepetit et al., 2013), triggering adjustments in protein accumulation and cellular function. Such longer-term and higher-order control process allows photosynthesis to switch between distinct self-maintenance regimes and corresponds to “regulation” rather than “dynamic stability” (Bich et al., 2016). Furthermore, the reciprocal influence between dynamic stability and regulation makes the interplay between external conditions, photosynthesis, and genome-wide regulation uniquely challenging to decipher. The aim of this paper is to investigate the parallel operation of (and interplay between) these two types of response in the context of the high-light response of the diatom *Phaeodactylum tricornutum*.

Exposure to higher light intensity immediately increases photon absorption by the antenna of both PSI and PSII, potentially exceeding photosynthetic capacity. When surpassing this threshold, the photosynthetic electron transport chain (ETC) becomes excessively reduced upstream of the rate-limiting cytochrome (cyt.) *b*_6_*f* complex, particularly at the secondary quinone acceptor Q_A_ of PSII. The over-reduction of Q_A_ is a critical step in the onset of light-induced damage. It directly promotes charge recombination within PSII, generating the formation of unstable chlorophyll triplet states. Crucially, chlorophyll triplets are susceptible to producing reactive oxygen species (ROS) which can cause photodamages to both PSI and PSII and, ultimately, photoinhibition (Barbato et al., 2020; Campbell and Serôdio, 2020; Tiwari et al., 2016). Photoinhibition induced by high light stress involves the photo-oxidation of PSII core PsbA and PsbB proteins in green organisms (Kale et al., 2017; Telfer et al., 1999) and results in a decline in PSII photochemistry, while damaged PSII act as non-photochemical quenchers (qI) (Nawrocki et al., 2021). Understanding how “dynamic stability” modulates this chain reaction across taxa and depending on environmental conditions remains a major frontier in photosynthesis research. This is particularly true in marine groups like diatoms, where research still lags behind that in plants, despite the equal contribution of terrestrial and marine compartments to global primary production (Field et al., 1998).

Among mechanisms granting “dynamic stability” to photosynthesis, the fast component of NPQ plays a central role in limiting ROS formation in PSII by dissipating excess excitation energy as heat (Goss and Lepetit, 2015). In *P. tricornutum*, NPQ is unusually simple compared to its compounded nature in other organisms (Short et al., 2023; van Amerongen and Croce, 2025), in which it often overlaps with state transitions that are inexistent in diatoms (Lepetit et al., 2022). It relies on two molecular actors: **i)** various isoforms of light-harvesting complex proteins — Lhcx and **ii)** the de-epoxidized xanthophyll-cycle pigment diatoxanthin (DT). As such, rapidly reversible NPQ in pennate diatoms consists of a single component, qZ, whose level is strictly proportional to DT interacting with Lhcx (Croteau et al., 2025). Unlike with PsbS or LHCSR3 and the NPQ/qE component in green organisms (Li et al., 2000; Peers et al., 2009), lumen acidification does not directly modulate Lhcx properties and NPQ/qZ in *P. tricornutum* (Buck et al., 2021; Giovagnetti et al., 2022). Instead, the xanthophyll-cycle enzymes determining DT concentration are coupled to photosynthetic outputs such as the transthylakoidal proton gradient and NADPH (Giossi et al., 2025; Goss et al., 2006). While Lhcx1 is constitutive, its abundance - as well as those of the inducible isoforms Lhcx2 and Lhcx3 - is regulated on a photoacclimative timescale (Bailleul et al., 2010b; Buck et al., 2021, 2019; Lepetit et al., 2013; Nymark et al., 2009; Taddei et al., 2016). Abundance of Lhcx isoforms and the size of the xanthophyll pool are regulated in response to growth conditions, which the cells detect via different sensors of external abiotic factors (Griffin and Toledo-Ortiz, 2022; Jaubert et al., 2023) and internal photosynthetic signals, such as the redox state of the plastoquinone (PQ) pool (Lepetit et al., 2013). Thus, while gene regulation determines the latent potential of NPQ/qZ (designing fast reversible NPQ in diatoms from here onward), its realized level under a given stress is the result of coordinated actions between two hierarchical responses.

Beyond NPQ, an intricate network of alternative electron fluxes contributes to photosynthetic resilience (Croteau et al., 2024), among which the cyclic electron flux (CEF) around PSI is of high interest. Crucially, CEF engages the cyt. b*_6_f* to contribute additional proton pumping into the lumen and thereby supports extra ATP production. In plants, NPQ is hypothesized to support tolerance to high-light stress through two linked effects: it alleviates excitation pressure on PSII, making room for increased CEF to boost ATP output (Li et al., 2018). Notably, this would link NPQ to a further layer of dynamic stability by influencing the interplay between PsbA degradation and its ATP-consuming repair cycle (Murata and Nishiyama, 2018). While this is vital in plants (Munekage et al., 2004), the CEF constitutive rate appears low in diatoms (Bailleul et al., 2015). This work investigates how NPQ/qZ shapes the “dynamic stability” of photosynthesis — focusing on its interaction with cyclic electron flow (CEF) and the PSII damages repair cycle — and examines whether NPQ/qZ also influences “regulations” mediated by changes in gene expression. Our reasoning was as follows: light-induced transcriptomic responses arise from the coordinated action of multiple sensors and signaling pathways that detect changes in the light environment, either directly (e.g. photoreceptors) or indirectly through shifts in the physiological state of the photosynthetic apparatus (e.g. retrograde signals). If NPQ/qZ actively modifies this physiological state, then NPQ/qZ could, in principle, also affect the transcriptomic responses triggered by high light.

To test this hypothesis, we made use of an experimental “NPQ-dial**”** created through a set of targeted genetically modified strains of the diatom *P. tricornutum* (Croteau et al., 2025; Giovagnetti et al., 2022). These transgenic lines accumulate different, yet stable, levels of Lhcx1 even under non-stressful low-light conditions, where NPQ/qZ is inactive (Blommaert et al., 2021; Croteau et al., 2025). As a result, all strains begin experiments from an indistinguishable physiological baseline, but once exposed to high light, their NPQ/qZ levels scale predictably with Lhcx1 abundance - providing a finely graded NPQ-dial spanning up to ten strains. This high-resolution gradient allowed us to establish phenomenological relationships between NPQ/qZ and multiple photosynthetic indicators, including Q_A_ redox-state, CEF activity, PSII damage/repair dynamics, and genome-wide transcriptional remodelling. In doing so, we found that nearly half of *P. tricornutum*’s transcriptome is modulated indirectly by NPQ/qZ-driven shifts in physiological state under light-stress, distinct from genes responding strictly to absolute light intensity and from stable housekeeping genes. This approach enables an integrated analysis of both the dynamic stability and regulatory responses of photosynthesis under high-light stress in a single coherent framework and provides a new lens for understanding holistic regulation in biological systems.

## Results

### Effect of NPQ/qZ on photosystems energy partitioning and redox state of intersystem chain

Ten *P. tricornutum* strains, including the wildtype (WT (Pt2)), an Lhcx1-KO and eight complemented strains from the latter, expressing ranging baseline levels of Lhcx1 (see Methods), were grown under a weak light intensity corresponding to the light-limiting regime of photosynthesis. Under these conditions, none of the strains trigger NPQ/qZ, which ensures that all strains share a roughly equal initial state as shown in (Croteau et al., 2025). The 10 strains were then exposed to a stressful light intensity of 570 µmol photons m^-2^ s^-1^, chosen to maximize contrasts in NPQ/qZ while reaching light-saturated regime for all strains (Supplementary Fig. S1). Once a steady-state was reached (≈5 min), we characterized the short-term functional response of photosynthesis.

We measured energy partitioning in the two photosystems and the redox state of intermediary transporters (Fig. 1). Like in our previous findings (Croteau et al., 2025), NPQ was linearly correlated to Lhcx1 abundance under saturating light exposure (Fig. 1a). This robust relationship implies that Lhcx1 abundance in each sample can be estimated non-invasively from maximal NPQ/qZ capacity from here onward, defining the *x*-axis and the colour scale. As a function of Lhcx1 amount across strains, the PSII yield, ϕPSII, decreased only slightly (Fig. 1e). Consequently, the increase in the yield of NPQ, ϕNPQ, from nearly 0 in Lhcx1-Ko to ≈ 0.6 in the strains displaying the highest Lhcx1 expression, is compensated by a roughly equivalent decrease in the yield of non-regulated heat losses, ϕNO (Fig. 1f). Therefore, steady-state PSII activity remains largely unaffected by NPQ level. The fact that PSII yield — and thus electron transfer — stays independent of NPQ at steady-state across all light intensity (see ϕPSII in Fig. S1) may appear counter-intuitive. However, it reflects the finely tuned NPQ/qZ response of *P. tricornutum*. Under light-limiting conditions, ϕPSII is similar in all strains because NPQ remains unengaged in all of them. Under saturating light, linear electron flux (LEF) becomes constrained by downstream, non photochemical processes — such as the cyt. b*_6_f* turnover or the Calvin cycle (Sukenik et al., 1987) — and is therefore largely decoupled from photochemical regulation, including NPQ.

**Fig. 1.**
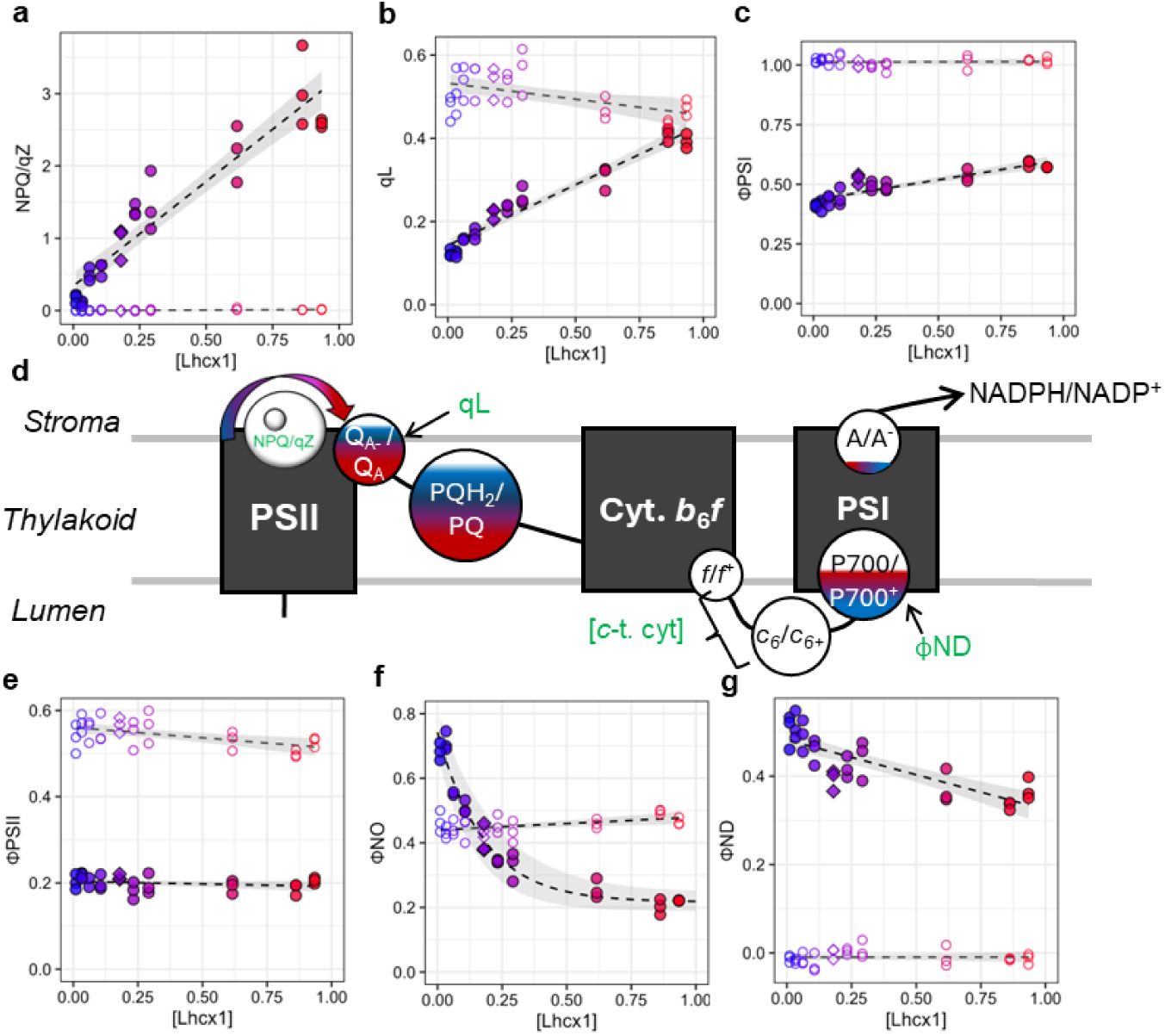
Lhcx1 abundance modulates the entire electron transport chain bioenergetics under high light in *Phaeodactylum tricornutum*. Diverse photophysiological parameters expressed as a function of Lhcx1 relative concentration (in r.u.) in 10 strains (Pt2-WT shown as diamond symbols) spanning a gradient generated via knockout and complementation, indicated by the colour scale. Each parameter was measured under steady-state low light (LL; 40 µmol photons m^-2^ s^-1^; open symbols) and high light (HL, 570 µmol photons m^-2^ s^-1^, closed symbols). **Top row**: NPQ/qZ (a), qL (which approximates Q_A_ redox state in the lake model (Kramer et al., 2004)), and thus, a proxy of the plastoquinone pool (PQ/PQH_2_) redox state) (b) and the quantum yield of photosystem (PS) I (ϕPSI) (c). Middle row: schematic representation of linear electron flow (black arrow) and the redox state of the photosynthetic electron transport chain under HL (d) – with more filled circles indicating more reduced electron carriers — coloured by Lhcx1 abundance. PSI acceptors (A/A^-^) reduction is calculated as 1-ϕPSI-ϕND and is consistently minimally reduced (Shown in Supplementary Fig. S1). **Bottom row**: ϕPSII (e), non-regulated heat dissipation by PSII (ϕNO) (f) and the yield of PSI donor-side limitation (ϕND, i.e. the proportion of oxidized PSI-P700) (g). Overall, the effects of Lhcx1 abundance (and thus NPQ/qZ) are minimal under LL, whereas under HL all parameters — except ϕPSII — follow a robust phenomenological relationship with Lhcx1. Triplicate per strain were measured, except one strain under LL (*n*=2), with the grey shading denoting 95% confidence intervals and fits statistical summaries found in Supplementary Fig. 2. The interactive effects between Lhcx1 concentration and light intensity (tested in three strains) are shown in Supplementary Fig. S1, which confirmed full oxidation of c-type cytochromes (c-*t* cyt.) under HL irrespective of Lhcx1 concentration (see Supplementary Fig. S3).

The PSII “lake model” approximates exceptionally well energy partitioning in pennate diatoms (Croteau et al., 2025), thus, the qL parameter represents a reliable proxy for the fraction of open PSII reaction centres, i.e. those with oxidized Q_A_ (Kramer et al., 2004). Under high light, qL increased markedly across the Lhcx1 gradient, indicating that the proportion of oxidized Q_A_ varied by roughly fourfold (Fig. 1b). Considering measurements were realized in steady-state, a parallel dependency of the redox state of the plastoquinone/plastoquinol (PQ/PQH_2_) upon Lhcx1 abundance is expected. Combined with the similar PSII yield observed among strains, this suggests that when NPQ/qZ increases while all other factors remain comparable, the NPQ/qZ-mediated shortening of PSII exciton lifetime is offset by a greater Q_A_ oxidation. As a result, LEF is largely unaffected across the NPQ/qZ gradient. In other words, when excitation pressure on PSII decreases, but electron fluxes towards plastoquinone is constant, Q_A_ becomes more oxidized. We also investigated how Lhcx1 amount affects the redox state of the photosynthetic electron transfer chain downstream of the PQ-pool. Unexpectedly, the special chlorophyll pair of PSI, P700, became more reduced with more Lhcx1 (see Methods; approximated by ϕND in Fig. 1g), whereas its acceptor-side limitation was low and invariant across strains (Supplementary Fig. S1). Therefore, the quantum yield of PSI increased with Lhcx1 abundance (Fig. 1c). Moreover, we confirmed that *c*-type cytochromes (*c*-t. cyt.), which comprise cytochrome *f* and cytochromes *c*_6,_ were almost fully oxidized at 570 µmol photons m^-2^ s^-1^ in all strains (Supplementary Fig. S3) in agreement with their lower redox potential relative to the P700/P700^+^ couple.

### Effect of NPQ/qZ on partitioning between linear and cyclic electron flows

Because PSII contributes only to linear electron flow whereas PSI supports both LEF and CEF, the positive change in PSI photochemical yield, relative to the negative change in PSII photochemical yield we report in (Fig. 1), suggests an increase in CEF activity. The intermediate increase of the photochemical rate (sum of both PS output) with NPQ/qZ (Fig. 2a and Supplementary Fig. S4) corroborates this hypothesis, although measuring absolute CEF values is notoriously challenging. Strictly speaking, it requires measuring simultaneously the absorption cross-section of both PS in parallel of their yields (Finazzi and Forti, 2004). However, since diatoms do not undergo state transition (Lepetit et al., 2022; Owens, 1986), the absorption cross-sections of both photosystems are not susceptible to change during the short-term experiments used here. This biological characteristic allowed the introduction of a new parameter, Δ(TEF/LEF), to estimate relative variations in CEF (see Methods). Here, TEF represents the total electron flow, i.e. the sum of CEF and LEF. This parameter is particularly advantageous because it only requires the measurement of PSI and PSII yields and is independent of the absolute values of their absorption cross-sections or of their stoichiometry (nonetheless reported in (Supplementary Fig. S5)), provided these remain constant throughout the experiment. The TEF/LEF ratio is equal to 1 in the absence of CEF and increases monotonically with increasing CEF (e.g. a value of 2 indicates equal rates of CEF and LEF). Thus, although Δ(TEF/LEF) is not a direct measure of CEF, it is a reliable indicator in species whose antenna size do not vary during the experiment. With this simplified approach we observed a linear relationship between the yield of energy dissipation via NPQ/qZ and Δ(TEF/LEF) under steady-state high light exposure (Fig. 2d).

**Fig. 2.**
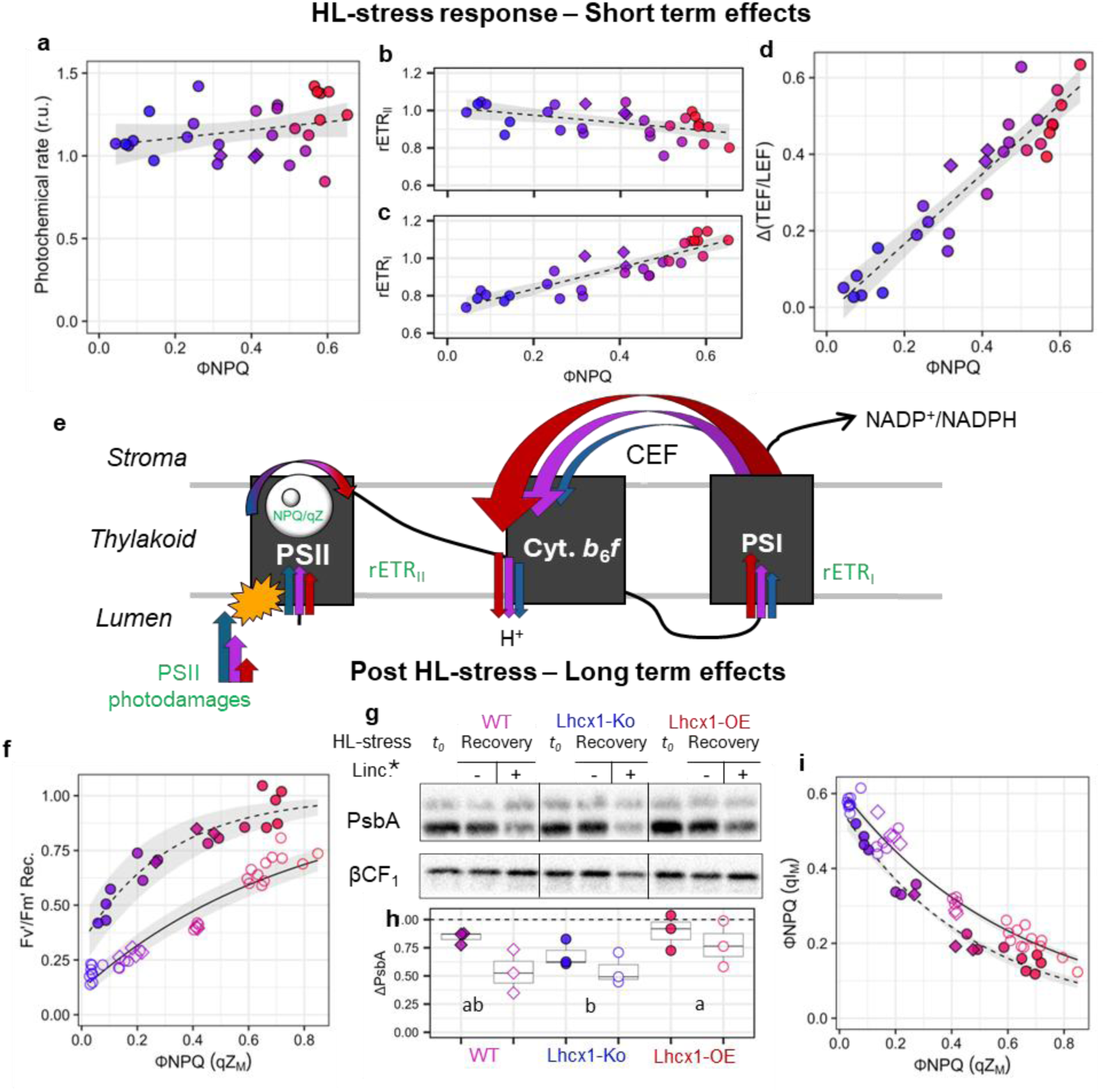
NPQ/qZ influences both PSI Cyclic electron flow (CEF) and PSII photoinhibition under high light (HL) in *Phaeodactylum tricornutum*. Data in panels a, b, c, d, f and i are plotted as a function of the PSII yield of energy dissipation by NPQ, ϕNPQ, also shown by the colour scale. **Top row**: Short-term (≈5 min) HL response (570 μmol photons m^-2^ s^-1^) across 10 strains spanning the Lhcx1 gradient (Pt2-WT shown as diamond symbols). The photochemical rate (representing the combined activities of PSI and PSII) (a), together with relative electron flow through PSII (rETR_II_) (b) and PSI (rETR_I_) (c), are normalized to the WT mean. From rETR_I_ and rETR_II_, HL-induced changes in CEF are estimated as the relative increase in the ratio between total electron flow and linear electron flow, Δ(TEF/LEF) (see Methods) (d). **Middle row**: Schematic illustration of how NPQ/qZ amplitude modulates CEF and PSII photoinhibition (e). **Bottom row**: Long-term (60 min) HL-response (600 μmol photons m^-2^ s^-1^). Recovered maximal PSII efficiency (*F_V_*’/*F_M_*’Rec.), after 1 h of HL-stress followed by 30 min of low light (15 μmol photons m^-2^ s^-1^), is plotted against the maximum contribution of the rapid NPQ component (qZ_M_) to ϕNPQ (illumination protocol in Supplmentary Fig. S7). This relationship was measured in control (closed symbols) and lincomycin (Linc.) treatments (average of 0.4 and 0.8 µg mL^-1^, fully blocking PSII repair, open symbols, see dose response in Supplementary Fig. S8) in six strains (f). Under the same conditions, immunoblots were performed on three strains (Lhcx1-KO strain, Pt2-WT and the Lhcx1-overexpressor (OE) LtpM) to quantify degradation of the core PSII protein PsbA, using ATP-synthase βCF1 as loading control (g). The significant effect of Linc. (*P* = 0.02) is indicated by * (g), whereas the letters in (h) denote a strain-specific trend considering the intrinsic variability of immunoblot measurements (*P* = 0.07, 2-way ANOVA, Tukey). Immunoblots material was sampled before HL stress (“no”) and after 30 min of low light recovery (“Recovery”) with (+) and without (-) addition of Linc. (0.4 µg mL^-1^), with changes relative to *t*_0_ in PsbA (ΔPsbA) in (h) and unedited blots in Supplementary Fig. S10. The maximum contribution of the slowly-relaxing photoinhibition-associated quenching component (qI_M_) to ϕNPQ is plotted against ϕNPQ (qZ_M_, see Supplementary Fig. S9 for the deconvolution of the two-components qZ and qI) in (i). Grey shading denotes 95% confidence intervals on the fits. The experimental data shown in panels a-d et f-i correspond to independent biological triplicates for all strains except one in a-d (*n*=2).

To further confirm the link between NPQ and CEF, we conducted a second experiment under light-limiting conditions. We monitored CEF during a transitory phase from saturating to low light, while NPQ/qZ relaxed from the distinct initial levels established in four strains during a preceding light-stress treatment (Supplementary Fig. S6). Under these conditions, PSII activity recovery was tightly coupled to NPQ/qZ relaxation over 5 min, while PSI activity almost instantly returned to a maximal steady value. This produced a similar relationship between Δ(TEF/LEF) and the yield of NPQ/qZ as the one observed under light-saturating regime (Fig. 2d). In both experiments, Δ(TEF/LEF) remained unchanged in the absence of NPQ (see Lhcx1-KO mutant and the *y*-intercept of the linear fit, Fig. 2d and Supplementary Fig. S6). This indicates that high light alone is not sufficient to directly trigger an increase in CEF, it requires the downregulation of PSII by NPQ/qZ. Crucially, CEF around PSI involves proton pumping across the thylakoid membranes. Accordingly, the increase in steady-state photochemical rate with Lhcx1 abundance, together with constant amplitude of proton motive force generated under illumination across strains (Supplementary Fig. S4), confirms that stronger NPQ/qZ coincides with faster proton efflux through the ATP-synthase, thereby augmented ATP production.

### Effect of NPQ/qZ on PSII photoinhibition

Based on the profound remodelling of photosynthesis under high light across our “NPQ-dial”, we examined the long-term effects of NPQ/qZ by monitoring PSII performance over a 1 h high-light stress, followed by 30 min under low light to allow for NPQ/qZ relaxation (see Methods and Supplementary Fig. S7). Six of the 10 strains, spanning the entire NPQ/qZ capacity range, were selected. To isolate NPQ/qZ relaxation from overlapping PSII repair, some samples were supplemented with lincomycin — a chloroplast translation inhibitor blocking PsbA turnover — at concentrations sufficient to fully abolish PSII repair (see intermediate dose-response in Supplementary Fig. S8) (Fig. 2f). All strains exhibited roughly equal PSII potential yield in the dark (*F_V_*/*F_M_*) before light stress, yet their recovered levels after low light exposure (*F_V_*’/*F_M_*’Rec.) varied two-fold under control treatment: Lhcx1-KO was 50% diminished, whereas high-Lhcx1 strains fully recovered (Fig. 2f). This indicates that PSII photodamages leading to photoinhibition scale inversely with NPQ/qZ. When PSII repair was inhibited, this *F_V_*’/*F_M_*’Rec. trend shifted downward (20 to 75% recovery) (Fig. 2f), reflecting the synergistic roles of PSII repair and NPQ/qZ to mitigate photoinhibition during and after high-light stress. As in green algae and plants, the loss of PSII photochemical potential is accompanied by a slowly relaxing form of NPQ, commonly referred to as photoinhibition-related quenching (qI). To quantify this phenomenon, we decomposed the biphasic relaxation of the NPQ yield under low light into its fast (qZ) and slow (qI) components (see Methods and Supplementary Fig. S9). As with *F_V_*’/*F_M_*’Rec., the level of qI detected at the end of low-light recovery could be expressed as phenomenological functions of the qZ levels reached during high-light stress (Fig. 2f, i). Immunoblotting of PsbA supported these observations, with greater loss in PsbA sampled at the end of the 30 min recovery detected in Lhcx1-KO than in the wildtype and an Lhcx1 overexpressing strain, and exacerbated losses in all strains when PSII repair was inhibited by lincomycin (Fig. 2g, h). While PsbA variations appeared less pronounced than fluorometric metrics, this is consistent with only a fraction of total PsbA being photochemically active, yielding more complex relationships between qI and *F_V_*’/*F_M_*’ (Lavaud et al., 2016).

Altogether, these results confirm that Lhcx1-dependent qZ protects PSII from photodamages and slows down PsbA proteolysis. Accordingly, a simple model of the compounded photoprotective effects of NPQ/qZ in diatoms can be proposed. Under saturating light, increased NPQ/qZ limits Q_A_ reduction, which in turn decreases the likelihood of charge recombination and photodamages, protecting PSII from photoinhibition. Additionally, Q_A_ and PQ-pool oxidation support enhanced CEF around PSI, increasing ATP production. This augmented energy supply sustains the ATP-consuming PSII repair mechanisms (Murata and Nishiyama, 2018), thereby multiplying the protective effect of NPQ/qZ under high-light stress.

### Half of P. tricornutum genome’s expression is affected by NPQ/qZ during high-light stress

Beyond enabling dissection of processes underlying photosynthesis “dynamic stability”, our NPQ-dial approach can be used to explore long-term regulation of photosynthesis and the broader cellular response to high light. Like other photosynthetic organisms, diatoms exhibit widespread transcriptional reprogramming under high light (Kan et al., 2023; Nymark et al., 2009; Truong et al., 2023; Zhou et al., 2022). We reasoned that the integrated signalling processes governing this gene-expression tuning could be divided into two distinct components: **i)** a direct response to *absolute* light properties (intensity and quality), expected to be independent of NPQ and potentially mediated by photoreceptors and, **ii)** an indirect response to *experienced* light, shaped by physiology downstream of photochemistry and therefore expected to be NPQ/qZ-dependent. Signals mediated by ROS or PQ-pool reduction (Escoubas et al., 1995; Jung et al., 2013; Lepetit et al., 2013) would fall into this category. Within this conceptual framework, the interplay between dynamic stability and longer-term transcriptional regulation can be readily examined. First, this allows us to differentiate genes directly responding to high light (NPQ/qZ-independent) from those indirectly regulated by NPQ/qZ-driven effects on photosynthesis. Second, genes of the former group can be classified according to their dependence on NPQ/qZ dynamics. To this end, we selected six strains spanning the full NPQ/qZ range and sampled transcriptomes at 0, 20, and 60 min of high-light exposure, for a total of 50 transcriptomes (Fig. 3a). Principal Component Analysis (PCA) revealed that nearly all of the variance is captured by two components: a rapid response which relaxed between 20 and 60 min (PC1, 61%), and a long-term acclimation pattern mostly emerging at 60 min (PC2, 30%) (Fig. 3b). Each component showed clear NPQ/qZ dependence at 20 min, which strengthened further after 60 min of high-light stress (Fig. 3c), indicating that rapid changes in photosynthetic physiology mediated by NPQ/qZ indeed propagate to changes in long-term gene regulation.

**Fig. 3.**
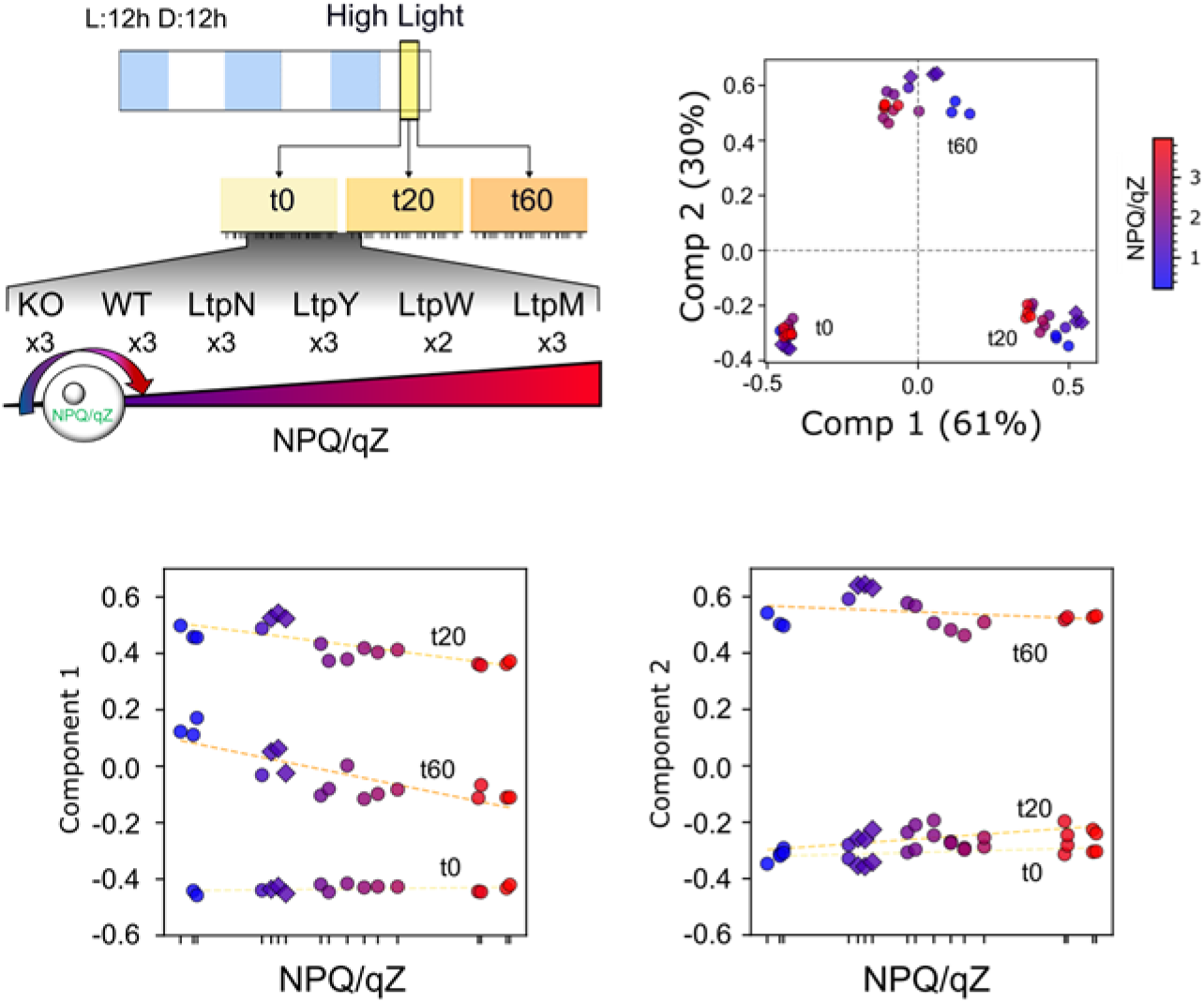
Experimental design of the transcriptomic response to high-light stress and variation visualization across an NPQ/qZ gradient. Six strains spanning the full gradient of NPQ/qZ capacity (colour scale) were transferred from growth light to a high-light stress (600 µmol photons m⁻² s⁻¹) and sampled at 0, 20 and 60 min for RNA extraction and transcriptome-wide sequencing (a). A principal component analysis (PCA) of all transcriptomes is shown in (b), with 91% of the variance explained by the first two components. Component 1 and Component 2 are plotted against NPQ/qZ for each time point in (c-d). One sample’s material was lost during extraction, and one biological replicate was excluded due to a strain that reverted during culturing, leaving 50 transcriptomes for analysis. Read mapping statistics and quality-control metrics for all samples are provided in Supplementary Table S1.

To further examine interplay between direct light cue and NPQ/qZ, we aimed at clustering genes by their expression profiles in terms of both high-light stress duration and NPQ/qZ capacity (defined as the maximal NPQ/qZ value reached by a given strain, see Methods). First, the weight of these two dimensions was uniformized by their standard deviation (Fig. 4a and Methods). From here onward, gene expression profiles are shown as weighted counts spanning the 50 transcriptomes, organized into three discrete snapshots: 0, 20 and 60 min, from left to right, and ranked from replicates with the lowest to the highest NPQ/qZ capacity within each panel (Fig. 4a). Of the 10,381 genes detected by RNA sequencing, 8,710 were retained for data transformation (see Methods and mapping in Supplementary Table S1) prior to K-means clusterization. The optimal cluster number determined by Silhouette (Supplementary Fig. S11) was as small as six (Fig. 4b). The content of each cluster was scrutinized by Gene Ontology (GO) enrichment analysis, revealing sharp contrasts in terms of Biological Processes and Cellular Component between clusters (Fig. 4c).

**Fig. 4.**
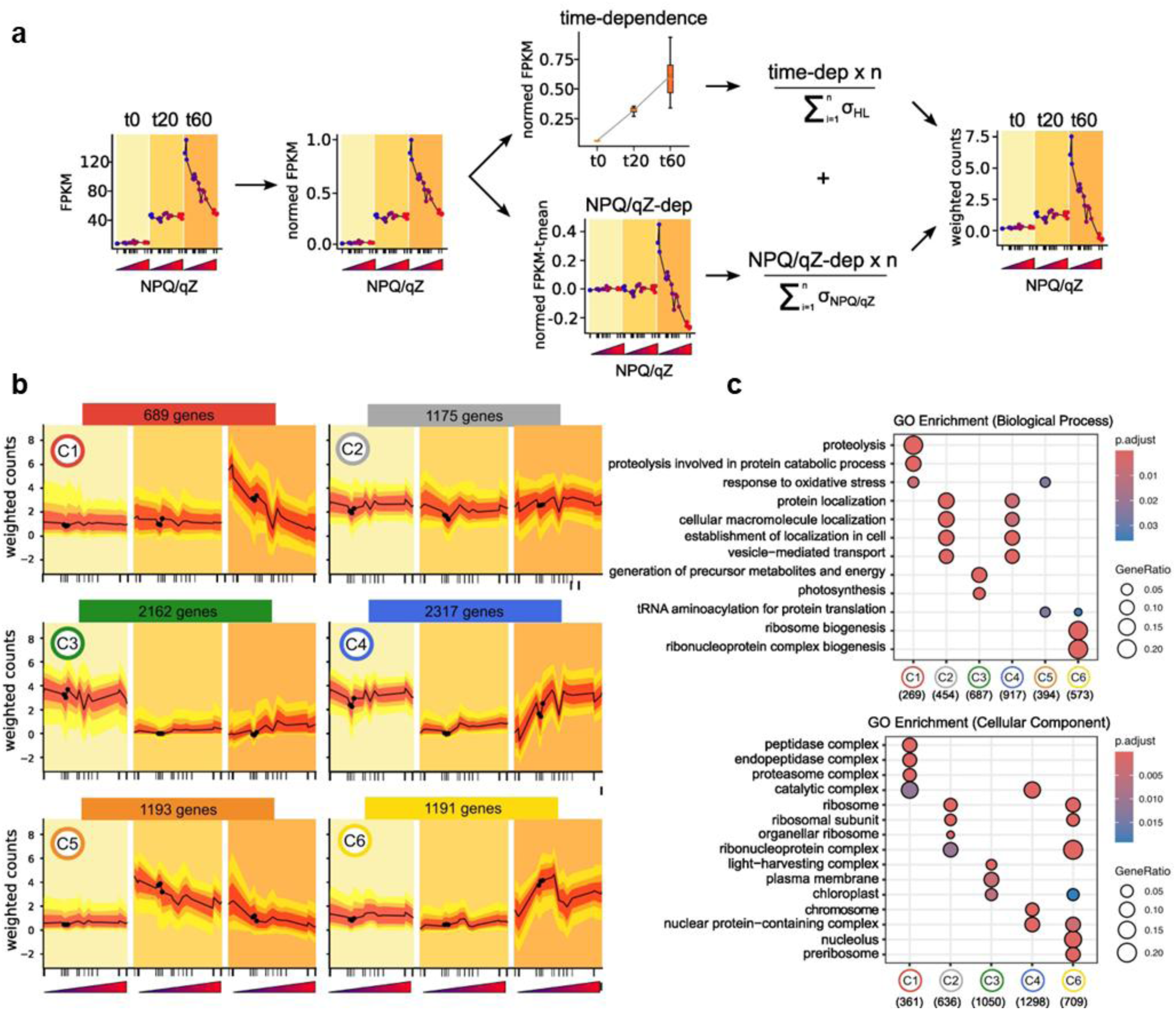
Genome-scale expression pattern clustering in response to high-light (HL) exposure time and NPQ/qZ in *Phaeodactylum tricornutum*. Example of the procedure of weighting of the two dimensions HL-exposure time and NPQ/qZ capacity — in the expression profile of a given gene (here CRTISO4), providing a plot of the “weighted counts” across the 50 transcriptomes (a). Clustering of 8,710 gene expression profiles into 6 main clusters by K-means (b) supported by the Silhouette analysis shown in Supplementary Fig. S11. Gene ontology (GO) significant enrichment for Biological Process and Cellular Component categories for each cluster is shown in (c); GO for Molecular Function is provided in Supplementary Fig. S12 and functional enrichment gene network of each cluster in Supplementary Fig. S13. In (b), black dots indicate the wildtype, while the yellow and red shadings denote dispersion across the gene profiles for all replicates within a cluster (the first–third quartiles and the 0.05–0.95 percentiles, respectively). Details about the weighting, gene selection criteria and clustering procedures are provided in Methods. All strains and replicates examined and their achieved NPQ/qZ during the experiment are found in Supplementary Table S1.

Among the six clusters (numbered arbitrarily by K-means), Cluster 2, which included 1,175 genes, showed the least variability across strains and over high-light exposure. It was largely enriched in genes associated with transport and structural functions, such as localization of key cellular constituents like proteins and macromolecules. Moreover, Cluster 2 was also tightly linked to ribosomal activity (Supplementary Fig. S12), hinting at a basal metabolic, or “housekeeping”, role for genes of this cluster (gene networks provided in Supplementary Fig. S13). By contrast, the 2,162 genes of Cluster 3 were strongly downregulated after 20 and 60 minutes of high-light stress, independently of strains’ NPQ/qZ levels. From GO enrichment analysis, Cluster 3 formed a distinctive “Photosynthesis regulation” gene set, with high GO ratios targeted to “light-harvesting complex” and “chloroplast” and involved in “photosynthesis” and “generation of precursor metabolites and energy” (Fig. 4c). In the gene network, a small sub-cluster associated with lipid catabolism was also visible in Cluster 3 (Supplementary Fig. S13c). The remaining four clusters pooled 5,390 genes whose expression profiles under high-light stress were markedly modulated in an NPQ/qZ-dependent manner. Two clusters included genes whose upregulation under high light was dampened by NPQ/qZ, either as early as 20 min (Cluster 5) or only after 60 min (Cluster 1). Cluster 5 was enriched in genes part of the oxidative stress response, but also in “tRNA aminoacylation for protein translation” (Fig. 4c). By contrast, on top of oxidative stress genes, the slower response of Cluster 1 was driven almost uniquely by genes associated with proteolysis, largely directed towards proteasomal and peptidase complexes. Cluster 4 included genes repressed at 20 min, but whose expression was restored at 60 min in an NPQ/qZ-dependent manner. Genes enriched in Cluster 4 were associated with similar overall biological processes as Cluster 2, but related to chromosomes and nuclear-protein complex, instead of ribosomes (Cluster 2). Additionally, many genes involved in key steps of the cell cycle, like mitosis and DNA replication, were greatly enriched in Cluster 4. Cluster 6 was induced exclusively after 60 min and showed a peculiar NPQ/qZ-dependent trend in weighted counts peaking around wildtype-like NPQ levels (Fig. 4b). An overwhelming fraction of Cluster 6 genes were involved in ribosome-genesis- and RNA-processing-related processes.

### NPQ-dependence of “regulation” of photosynthesis

To explore the ramifications of the observed interplay between the two responses in more detail, we focus on the expression profiles of key gene-sets and known biosynthetic pathways involved in the high-light response. These included nuclear-encoded and plastid-targeted genes that have already been reported for their differential responses to high-light cues (Agarwal et al., 2023; Kan et al., 2023; Nymark et al., 2009; Truong et al., 2023; Zhou et al., 2022). Specifically, we examined expression patterns of genes encoding enzymes of chlorophylls and carotenoids biosynthesis pathways, as well as fucoxanthin-Chl *a*/*c* binding proteins (FCP) from several subgroups: Lhcf (major antenna proteins), Lhcr (red algae homologs mostly associated to PSI), Lhcx and high-light-induced proteins (HLIP). Across these functional categories comprising a total of 109 genes, 90 were found among only three clusters: Cluster 3, corresponding to genes repressed by high light independently of NPQ/qZ, and Cluster 1 and 5, grouping genes induced by high light but counteracted by NPQ/qZ (Fig. 4). These two opposing dynamics exposed a regulatory split between genes defining photosystems’ light-harvesting-antenna arrangement and functionality. Genes responsible for increasing light absorption were predominantly repressed by high light regardless of NPQ/qZ levels (Cluster 3). This includes all Lhcf genes but Lhcf15, eight Lhcr genes and the dark-induced gene Lhcx4 (Nymark et al., 2013) (Fig 5a). Cluster 3 was also enriched in chlorophyll biosynthesis pathway genes — 21 out of 24 — including the recently identified chlorophyll *c* dioxygenase (Jiang et al., 2023) (Fig. 5c). Interestingly, GLU-RS2 was the only gene repressed by NPQ/qZ (Cluster 5). This gene is involved in the initial step of the pathway and plays a key role in supplying Glu-tRNA for tetrapyrrole (chlorophyll/heme) biosynthesis, in parallel to the canonical cytosolic Glu-RS1. For carotenoid and xanthophyll biosynthesis, 21 out of 44 genes also belonged to Cluster 3 (Fig. 5d).

**Fig. 5.**
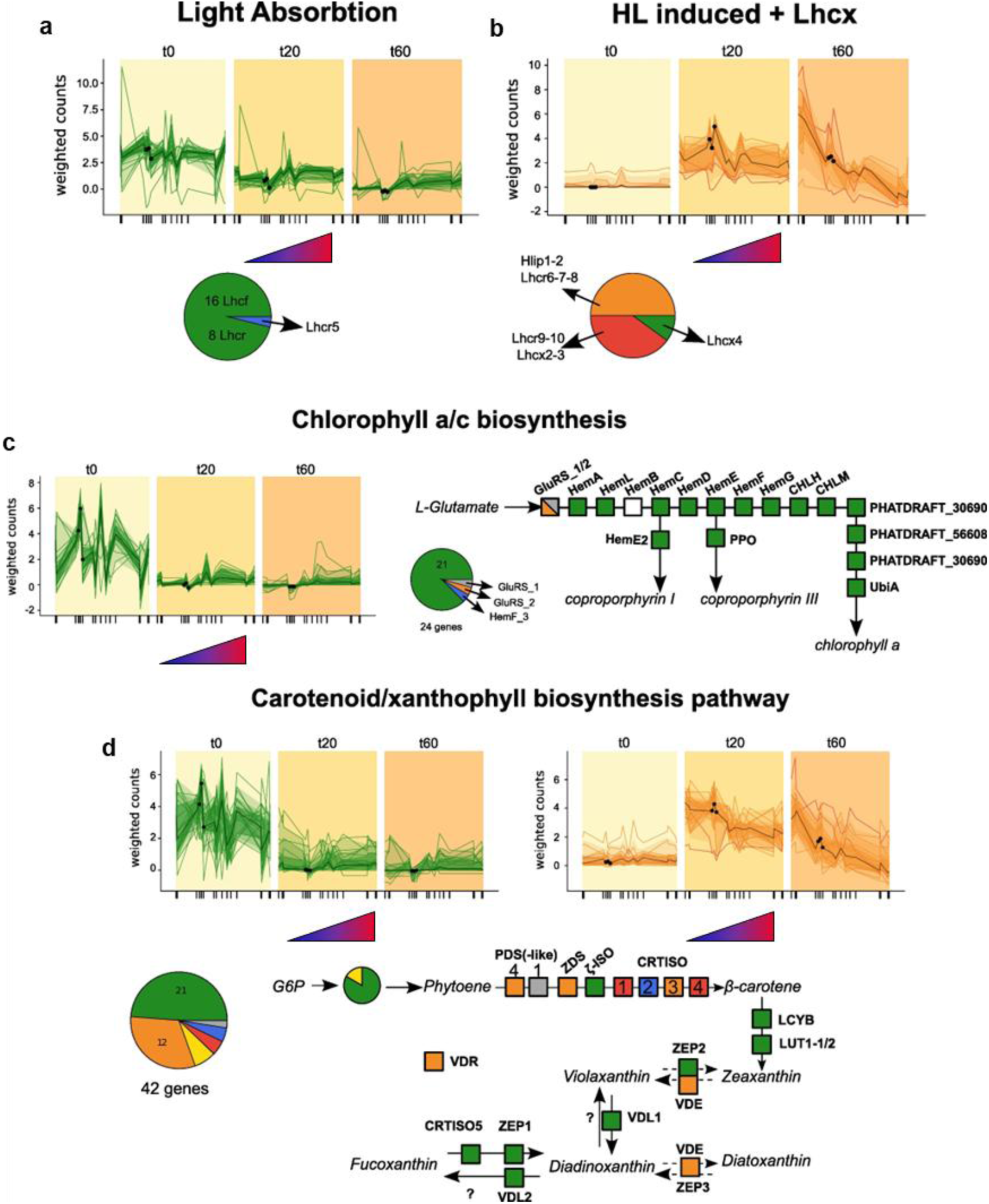
Cluster distribution of genes encoding antenna proteins and enzymes of the chlorophyll and carotenoid biosynthesis pathways. Each panel includes a pie chart indicating the proportion of genes assigned to the six expression profiles clusters (cluster colours as in Fig. 4) and representative expression profiles for genes belonging to Cluster 3 (green), 1 (red), and 5 (orange). (a) Light-harvesting complex (LHC) proteins in *Phaeodactylum tricornutum*, including all Lhcf (major antenna) and nine Lhcr (red-algal PSI homologs). (b) High-light–induced antenna proteins, comprising five Lhcr (as in Nymark et al., 2009) and three Hlip/Lhcx isoforms (except Lhcx1). (c) Chlorophyll *a* and *c* biosynthesis genes together with a simplified scheme of the chlorophyll biosynthetic pathway (white square because HemB is not identified by KEGG in *P. tricornutum*). (d) Carotenoid and xanthophyll biosynthesis genes and a simplified pathway based on Cao et al., (2023). The 13 genes involved in the conversion from glycerol-3-phosphate (G3P) to phytoene are shown only as a pie chart. Numbers above coloured squares indicate the number of isoforms; individual isoforms are labelled within squares when their cluster assignment differs. Dashed arrows denote xanthophyll cycles. VDE catalyses both violaxanthin to zeaxanthin and diadinoxanthin to diatoxanthin de-epoxidation; placement of ZEP3 follows Giossi et al., (2024). Question marks mark unknown intermediates or enzymes. The VDE-related protein (VDR), of unknown function, is shown outside the pathway.

The other side of the regulatory split reveals that the high-light-driven upregulation of genes involved in photoprotection — xanthophyll cycle enzymes, Lhcx and high-light-induced proteins (HLIP) — is further shaped by an NPQ/qZ-dependent regulation layer (Cluster 1 and 5). As such, their expression profiles are characterized by a high-light induction dampened by NPQ/qZ (Fig. 3 and 5b). Five Lhcr genes, previously reported to be unexpectedly induced under high light (Kan et al., 2023; Nymark et al., 2009), showed such NPQ/qZ-dependent expression profiles (Fig. 5d). Additionally, Cluster 1 and 5 contained HLIP and Lhcx2 and Lhcx3 isoforms, as well as 16 of the 44 carotenoid-related genes. Regarding the xanthophyll biosynthesis pathway (downstream of the upper carotenoid branch), the genes encoding VDE and ZEP3 — enzymes regulating the xanthophyll cycle and thus NPQ/qZ formation — were found in Cluster 5, together with the VDE-related gene (VDR), whose function remains unknown. By contrast, genes encoding all other enzymes with xanthophyll substrates were found in Cluster 3 (Fig. 5d). The distribution of xanthophyll biosynthesis genes between Cluster 3 and Cluster 5 is informative considering their dual roles in diatoms: enzymes proposed to lead to fucoxanthin synthesis (Cao et al., 2023) — which strongly contribute to light absorption — are downregulated in an NPQ-independent manner (Cluster 3), whereas xanthophyll-cycle enzymes involved in photoprotection (Giossi et al., 2025) are induced by high light, but this induction is mitigated by NPQ/qZ (Cluster 5).

### Holistic view on the cellular response to high light

Strikingly, the expression of nearly half of the genome (5,390 genes, Fig. 4) under high-light stress is fine-tuned in an NPQ/qZ-dependent manner. Our functional analyses (Fig. 1 and 2) show that, upon high-light exposure, cells conform to an NPQ/qZ-defined photophysiological state reflected in the redox status of PSII acceptors and PSI donors, the balance between CEF and LEF, which in turn, influence the extent of ROS-induced PSII photoinhibition on the long-term. When combined with genome-wide clustering of expression profiles, this functional framework provides mechanistic grounding for how short-term adjustments in photosynthesis shape long-term transcriptional regulation, potentially via retrograde signalling pathways. Figure 6 integrates these observations by comparing the temporal dynamics of two extreme scenarios — cells exhibiting either low (<0.5) or high (>3) NPQ/qZ levels (shown in blue and red, respectively) — to provide a holistic perspective on the cellular response to high light.

**Fig. 6.**
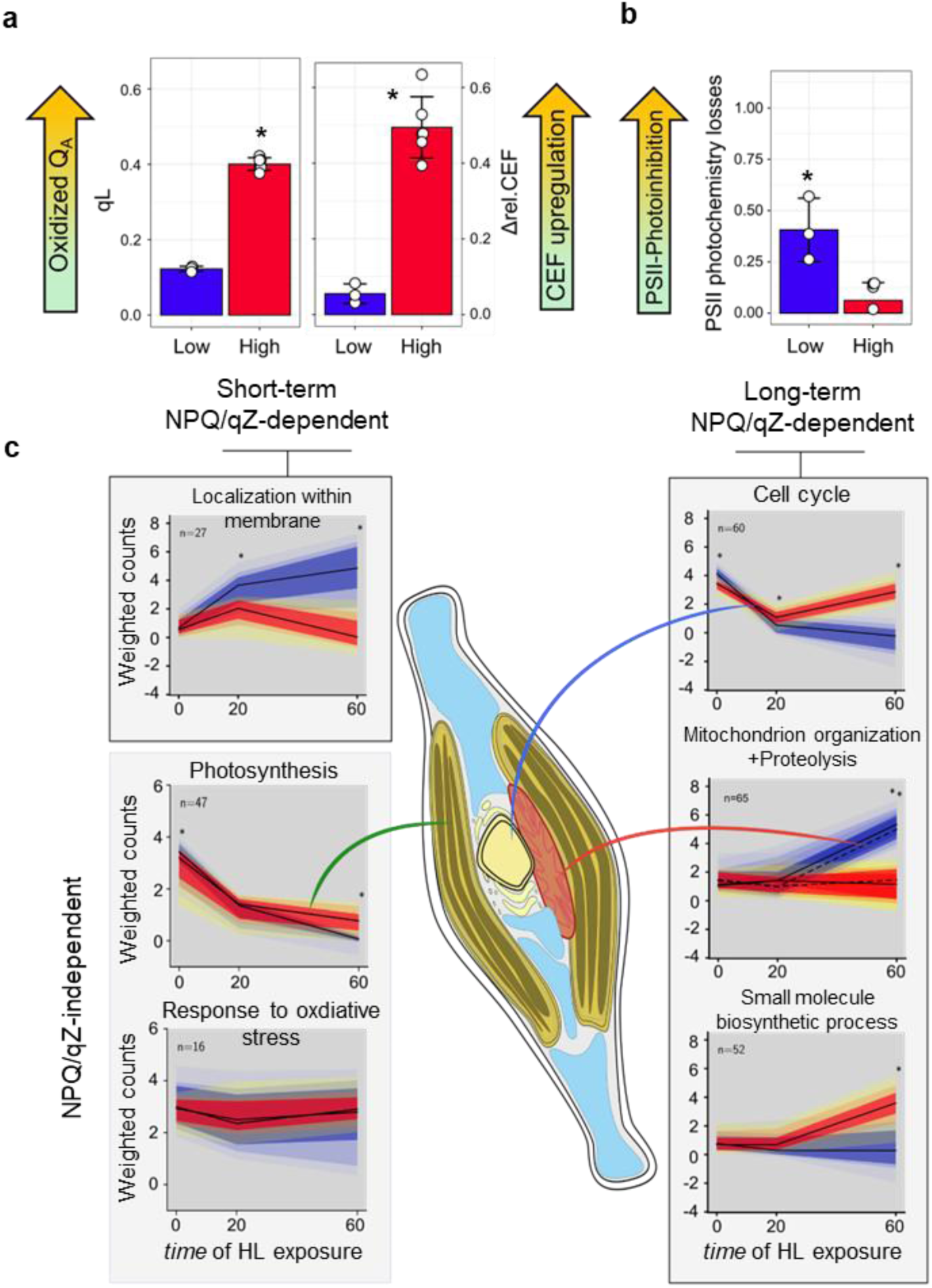
Holistic view of the response of “low-NPQ” (blue) and “high-NPQ” (red) cells to high light (HL). Short-term (a) and long-term (b) functional response to HL include variable reduction of Q_A_ and PQ-pool, induction of CEF around PSI and PSII photoinhibition. The time-series of weighted counts of seven GO terms (Mitochondrion organization and proteolysis (in dashed line) are shown in a same panel), representing typical biological processes associated to each of the six clusters in Fig. 4, are shown in (c), together with the cellular compartment associated to certain processes in the diatom cell scheme. Star (*) symbols indicate significantly higher value (independent *t*-test, *P.*value<0.05) at a given time and in (c), light and dark coloured shadings denote weighted counts dispersion for a given gene, for either low- or high-NPQ strains, across the first–third quartiles and the 0.05–0.95 percentiles, respectively.

The short-term transcriptomic response to high light was assessed at 20 min, coinciding with rapid functional adjustments observed in high-versus low-NPQ/qZ strains, including a less reduced Q_A_ and PQ-pool and enhanced relative CEF (Fig. 6). Only Cluster 3, 4, and 5 displayed differential expression relative to *t*_0_ at this early time point, forming two contrasting patterns: repression of genes involved in core cellular activity and induction of genes involved in the stress-response. The repressed category comprised genes associated with DNA replication, the cell cycle, and mitosis — predominantly in Cluster 4 — as well as photosynthesis genes in Cluster 3. If a transient inhibition of photosynthesis genes expression prevents further oxidative damages, a parallel halt in replication could limit the propagation of replication errors under stress, creating a safer window for cell-wide reprogramming proceeds. In both cases, this rapid downregulation was NPQ/qZ-independent. High-light induced genes fell largely within Cluster 5 and already showed a strong contrast between low- and high-NPQ/qZ strains at 20 min, reflected by “oxidative stress response” genes (Fig. 6c). This cluster contains photoprotection genes (Lhcx, HLIP, xanthophyll-cycle enzymes), redox-regulatory components — including several glutaredoxins (GLRX1, GLRXC1, GLRXC2, GLRXM1), thioredoxins (Trx-f, Trx-m, Trx-y), glutathione reductases (Gsr_1, Gsr_2) — as well as oxygen detoxification enzymes (e.g. SOD1, GPX2). Together, these observations support that photosynthetic signals — such as the PQ-pool redox state or ROS generation — may contribute to the regulated induction of these genes, while acknowledging that additional triggers are likely involved (see Discussion).

After 60 min, the more reduced Q_A_ observed in low NPQ/qZ strains is associated with marked PSII photoinhibition, potentially exacerbated by their inability to upregulate CEF and meet the ATP demands of the PSII repair cycle (Fig. 6b). This dual role of NPQ/qZ — limiting photodamage and supporting repair — is reflected in the behaviour of Cluster 5 stress-response genes, which had fully relaxed by 60 min only in the high NPQ/qZ strains. In low NPQ/qZ cells, the stress signature deepened with the pronounced induction of Cluster 1, which was strongly enriched in proteolysis related genes. Additionally, Cluster 1 included numerous genes involved in mitochondrial organization, consistent with a trade-off between upregulation of CEF (negligible in low NPQ/qZ cells) and chloroplast-mitochondrion coupling as an alternative energy outlet (Bailleul et al., 2015). Following the rapid shutdown of Clusters 3 and 4 at 20 min, genes linked to the cell cycle nearly returned to their initial expression levels after 60 min only in strains capable of high NPQ/qZ, whereas repression of photosynthetic genes persisted across all strains (Fig. 6c). This pattern suggests that, following high-light-induced arrest of the cell cycle, mandatory checkpoints required for growth resumption are more readily met when high NPQ/qZ mitigates the deleterious consequences of the stress. Meeting these checkpoints may depend on downstream metabolic reconfiguration that alleviates energy-flux imbalances. In contrast, sustained repression of photosynthetic genes is consistent with an early transcriptional signature of the cells’ long-term (day scale) photoacclimation trajectory toward growth under abundant light (Kan et al., 2023; Nymark et al., 2009).

Finally, Cluster 6 comprises genes involved in tRNA processing and ribosome biogenesis. By contrast with the invariant Cluster 2, also associated with ribosomal processes, this reveals two distinct ribosomal modules: a housekeeping set in Cluster 2 and a set induced after 60 min to support the re-engagement of translation and growth once photoprotection is established in Cluster 6. Combined with the recovery of cell-cycle-related genes in high NPQ-qZ strains, this supports that photoprotected cells are better positioned to resume growth and utilize the products of photosynthesis. Although the upstream sensors that trigger this metabolic transition remain unclear, they likely entail the integration of several signals, among which sufficient energy availability and controlled ROS levels could be essential. The coordinated behaviour of Cluster 1 and 6 — associated with proteolysis and protein synthesis, respectively — foreshadows profound divergences in proteome remodelling between low- and high-NPQ/qZ cells after 60 min.

## Discussion

Beyond relieving excitation pressure on PSII, NPQ/qZ operates within a feedback loop that provides dynamic stability to photosynthesis, making it more robust to perturbations. We dissected this functional chain reaction across a large set of mutants forming an experimental NPQ-dial, thanks to their wide-ranging Lhcx1 abundances. Critically, NPQ was fully inactive under growth conditions. Therefore, for all experiments, all strains started from equal initial states despite pronounced contrasts in their latent NPQ/qZ capacities. This shared baseline made the combination of functional characterization and whole-genome transcriptomic analysis uniquely powerful for probing the interplay between NPQ/qZ-mediated dynamic stability and slower regulation of gene expression. Our central hypothesis was that because NPQ extent determines the state into which the photosynthetic apparatus conforms under high light stress, it should indirectly modulate the available endogenous cues to the cell for integrating information and thus influence gene expression. We indeed observed that a large fraction of the genome — roughly 50% — was regulated in an NPQ/qZ-dependent manner during high-light stress. To reach this conclusion, gene expression was resolved across two dimensions: a direct response to *absolute* light intensity and an indirect one, reflecting light *experienced* by the cell for a given NPQ/qZ level. This framework bridges two traditionally distinct approaches in biology: reductionism, which analyses isolated components (in this case, NPQ), and systems biology, which seeks to understand emergent properties arising from complex interactions (such as genome-wide transcriptional responses). Integrating the strengths of each perspective remains a major challenge and often relies on computationally heavy tools such as genome-scale modelling and hybrid modelling. Our strategy offers a complementary route by “pulling the thread” of a unique and well-defined photosynthetic feedback loop — NPQ/qZ — as an entry point to expose systemic consequences arising from a perturbation by light stress. This strategic approach yields a comprehensive map of the high-light response in a diatom cell, in which diverse photosynthetic and transcriptional responses are described as functions of empirically measured NPQ/qZ. In doing so, it merges mechanistic insights with practical experimental simplicity.

The fingerprints of NPQ/qZ were first identified by examining the redox state of electron carriers from Q_A_ to the PSI acceptor-side (Fig. 1). While NPQ is generally assumed to favour oxidation of Q_A_ and the PQ-pool, our data also highlight a less intuitive fact: more oxidized Q_A_ is not necessarily accompanied by an increase in the rate of LEF. Additionally, the redox state of the PSI donors were strongly reduced as NPQ/qZ increased, while no changes in the redox state of PSI acceptors was found, leading to an unexpected increase in PSI electron flow proportional to NPQ/qZ. Together, these effects are consistent with an NPQ-dependent induction of CEF during high-light exposure. Importantly, CEF remained unchanged in the absence of NPQ, as seen in the Lhcx1-KO mutant under both saturating and limiting irradiance (Fig. 2d, Supplementary Fig. S6). This demonstrates that CEF activation is linked to the *experienced* light intensity — set by NPQ-driven downregulation of PSII — rather than to the absolute photon flux.

In diatoms, CEF is typically considered low under non-stressful conditions (Bailleul et al., 2015), but it could play a crucial role in overcoming nutrient starvation (Thamatrakoln et al., 2013), anoxia (Gain et al., 2023), and cold environments (Goldman et al., 2015). Our results extend this list to include high-light stress, provided that it induces NPQ/qZ. Strikingly, CEF does not merely compensate for the modest NPQ-induced decrease in LEF: the increase in PSI yields exceeds the relative decrease in PSII yields when both are represented as a function of NPQ/qZ (Fig. 2). Thus, CEF appears actively upregulated, which is further corroborated by the reduction of PSI donors with increasing NPQ/qZ. Yet, even with the measured faster photochemical rate (Fig. 2a), derived from a completely independent observable, this “overcompensation” by CEF is puzzling under a light-saturating regime. While the molecular components of CEF remain unidentified in diatoms (Croteau et al., 2024), and the competition dynamics between LEF and CEF are poorly understood in any organism, two classical bifurcation points are known: PSI acceptors (ferredoxin or NADPH) and the PQ-pool (Alric, 2015). In our framework, PSI acceptor-side limitation remained constant regardless of NPQ/qZ levels (Supplementary Fig. S1). This positions the redox poise of the PQ-pool — closely reflected by the relationship between qL and NPQ/qZ — as the most likely dominant factor controlling CEF-to-LEF ratio in diatoms.

Turning our functional analysis to longer-term consequences, we found that PSII photoinhibition following a 1 h light stress can be expressed as a function of NPQ/qZ, , whether quantified as the decrease in recovered *F_V_*’/*F_M_*’ or the extent of persisting qI quenching (Fig. 2). Greater losses in PSII-core protein PsbA proteins in Lhcx1-Ko, and when lincomycin is used, support that photodamages contribute to compromised PSII photochemistry. Transcriptomic analysis further strengthens this scheme with proteolysis genes being strongly overexpressed in the absence of NPQ/qZ after 60 min of high-light stress (Cluster 1, Fig. 4). Interestingly, the gene encoding the FtsH protease (Campbell et al., 2013), responsible for unfolding and degrading PsbA in photoinactivated PSII, was also overexpressed at low NPQ/qZ, but induced more rapidly (Cluster 5, Fig. 4). Overall, our data support the hypothesis that NPQ/qZ plays a dual role in preventing not only photodamages to PSII, but a cell-wide oxidative stress. First, it reduces excitation pressure on PSII, which limits excessive Q_A_ reduction, thereby reducing the risk of charge recombination and ROS production, which can damage a wide range of macromolecules (Pancheri et al., 2024; Ramel et al., 2012). Second, it stimulates CEF, which enhanced ATP production, essential for PSII repair (Murata and Nishiyama, 2018) and likely for other ATP-intensive processes part of the stress response, consistent with the numerous AAA+ATPase domains found encoded by Cluster 1 genes.

Among the processes we describe as functions of light exposure and NPQ/qZ, several have established roles as photosynthetic signals controlling gene expression in *Arabidopsis* and other model systems (Chan et al., 2016). This offers a novel avenue to investigate crosstalk between functional photosynthesis and transcriptomic regulation, with retrograde signalling pathways potentially bridging the two, a mechanism for which knowledge is currently sparse in diatoms. The redox state of the PQ-pool is one of the best established photosynthetic cues involved in retrograde signalling (Escoubas et al., 1995; Pfannschmidt et al., 1999). In *Arabidopsis*, several hundred Plastid Redox-Associated Nuclear Genes (PRANGs) respond to diverse signalling cascades originating from the PQ-pool redox state (Jung et al., 2013). In diatoms, a more reduced PQ-pool (manipulated via light and chemical manipulation) has been linked to the induction of Lhcx2-3 (Lepetit et al., 2013), fully consistent with the Cluster 5 membership of these isoforms in our data. Moreover, in *P. tricornutum*, translation of Lhcx2-3 can be further accelerated by an Aureochrome 1c pathway (Zhang et al., 2024), or repressed by a cryptochrome/photolyase pathway (Coesel et al., 2009), two photoreceptors that directly sense light. This illustrates that even genes whose expression depends directly on light cues can be further modulated by NPQ-driven effects, helping explain why, at the phenomenological level, such a large proportion of gene profiles appear regulated by *experienced*, rather than *absolute*, light intensity. In the future, combining our NPQ-dial with the usual chemical treatment approach could be instrumental in untangling complex interacting pathways and fast-tracking the assembly of a diatom-specific PRANG atlas.

Under light stress, increasing ROS generation provides the cell with additional information guiding gene expression (Foyer, 2018; Laloi and Havaux, 2015). Our results reinforce the view that by favouring the oxidation of Q_A_ (and the PQ-pool by extension), higher NPQ/qZ mitigates the formation of chlorophyll triplet states and thereby reduces singlet oxygen production and its damaging effects on PsbA (Telfer et al., 1999). Because of its extremely short lifetime, the direct actions of singlet oxygen are limited to the vicinity of PSII, and its signalling functions typically operate indirectly. Such indirect pathways occur via plastidial intermediates, such as EXECUTER proteins in *Arabidopsis* (Lee et al., 2007), or more broadly via oxidized carotenoids and lipid by-products (Pancheri et al., 2024; Ramel et al., 2012). The exacerbated PsbA degradation together with the extensive proteolysis genes induction in the Lhcx1-Ko provide novel experimental support for the existence of such macromolecule-degradation signals in diatoms. Alternatively, H_2_O_2_, a more stable ROS associated with PSI acceptor-side limitation and the Mehler reaction, can diffuse further within the plastid and activate multiple signalling cascades (Chan et al., 2016; Foyer, 2018). Many redox-regulatory components that are induced by exogenous H_2_O_2_ in diatoms (Rosenwasser et al., 2014) were affiliated with Cluster 5 in our dataset (listed in Results). Yet our functional analysis suggests that PSI acceptor-side limitation was low across strains, independently of NPQ/qZ levels (Supplementary Fig. S2). Notably, the stromal glutathione-ascorbate redox hub plays vital signalling functions, including in diatoms (Volpert et al., 2018), and often independently of ROS production (Foyer and Noctor, 2011). Speculatively, we propose that Lhcx1-Ko’s inability to enhance CEF as an electron-recycling route, could shift the glutathione-ascorbate redox poise, thereby activating genes that are sensitive to either glutathione-ascorbate or H_2_O_2_-mediated pathways.

Finally, this work provides a comprehensive dissection of a crucial functional feedback loop in diatoms photosynthesis, revealing intricate interactions between NPQ/qZ, cyclic electron flow and PSII photodamages. The Lhcx1 titration will continue to provide an ideal framework to expand this map of diatoms’ dynamic stability to other facets of their light-stress-response, such as alternative electron fluxes (Burlacot, 2023), or in combination with other system-wide methods, like targeted metabolomics (Zhou et al., 2022). Additionally, the gene clusters uncovered here, each organised around a coherent physiological theme, provide strategic subsets for seeking uncharacterized genes crucial to diatoms’ light-stress-responses in the future. Yet the broader impact of this work may lie less in the specific findings than in the conceptual framework it establishes, providing a transferable blueprint for studying regulators in their native dynamic environment. Instead of stripping a dynamic module from its physiological context, this approach engineers only its latent capacity, enabling its observation as an embedded node within a system upon perturbation. By preserving the natural regulatory architecture, the method moves beyond comparison of isolated actors to reveal the conserved feedback motifs that underpin photosynthetic resilience.

## Material and Methods

### Culture and growth conditions

Wildtype *P. tricornutum* Bohlin CCAP 1052/1A (Pt2) and 9 transgenic Lhcx1 lines previously characterized (Croteau et al., 2025) were grown in *f*/2 medium prepared from filtered (0.2 µm) seawater, in Erlenmeyers of 250 or 1000 mL depending on the volume required for a given set of experiments. In summary, the transgenic lines employed in this study consist of the Lhcx1 knock-out (Ko) strain (Ko6) together with eight of the twelve complemented lines described in Croteau et al., (2025). These complemented lines, which express varying levels of Lhcx1 in the Ko6 background, were originally generated by (Giovagnetti et al., 2022). Depending on the NPQ resolution required to address each experimental question, and on the practical constraints associated with sample volume and processing time, different numbers of strains were used. For example, 10 strains were included to characterize the steady-state bioenergetics of the electron transport chain, whereas three strains were used for the photoinhibition experiment conducted on large culture volumes to allow PsbA quantification by immunoblotting. In all cases, the selected strains spanned the broadest possible range of NPQ/qZ capacities (see below). Cells were grown under 5 µmol photons m^-2^ s^-1^ and 12: 12 photoperiod, those light conditions ensuring the basal expression of only the Lhcx1 isoforms (Croteau et al., 2025). Temperature was maintained at 19^°^C for growth and all experiments. Cultures cell density was monitored every 3 days for cell concentration using a Multisizer 3 Coulter counter (Beckman Coulter, CA, USA) and dilution were performed to always maintain the cultures in exponential growth (0.7-to-1.5×10^6^ cells mL^-1^). All experiments began between 4 and 6 hours into the light phase of the photoperiod to improve reproducibility.

### Fluorescence and spectroscopic measurements

For the *steady-state light exposure* experiments, samples of 50 mL were concentrated 10-fold by centrifugation (5000 RPM, 6 min), supplemented with ≈10% w/v Ficoll (to slowdown cell sedimentation during measurements), and then left to recover under gentle agitation and very low light (approximately 1 µmol photons m^-2^ s^-1^) for at least 30 min. Experiments were conducted in a JTS-10 spectrophotometer (Biologic, France), allowing to measure Chl *a* variable fluorescence and absorption change signals at different wavelengths by equipping the detecting light with specific interference filters (3-8 nm bandwidth). Two absorption change signals were probed. First, the movement of electrons and protons across the thylakoid membrane generate an electric field which induces the electrochromic shift (ECS) of pigments absorption spectrum, whereby change in the electric potential gradient can be observed by probing the linear ECS (ECS_lin_) (Bailleul et al., 2010a). Second, changes in the redox state of electron carriers, namely P700 and *c*-type cytochromes (*c*-t cyt., sum of cyt. *f* and cyt. *c*_6_, replacing plastocyanin in diatoms), result in absorption changes in the green and far-red regions of the spectrum, respectively. The *c*-t cyt. and ECS_lin_ signals were obtained through measuring transient absorption changes at 520, 554 and 566 nm with BG39 filter on the measuring and reference photodiodes (Schott (Mainz, Germany)) as in (Bailleul et al., 2015). Signals of interest were then calculated as *c*-t cyt. = [554]-0.4*([520]+[566]), and ECS_lin_=[520]-0.25* *c*-t cyt. Absorption differences related to the redox changes of P700 were measured at 705 nm – 730 nm with a high pass RG695 filters on the measuring and reference photodiodes, the difference between the two wavelengths allowing to eliminate the scatter signals. For every biological replicate, a first sub-sample (1.3 mL) was used to measured absorption difference following a saturating single turnover flash from a dye laser (690 nm) allowing to determine the amplitude of the ECS_lin_ signal corresponding to a single charge separation per PS, and the oxidation of a single *c*-t cyt. per PSI. Those two values were used for normalization of ECS_lin_ and *c*-t cyt signals. The maximal absorption change signal corresponding to full oxidation of P700 (*P_M_*) was then measured at the end of a 20 ms saturating pulse (red light, 660 nm, 6000 µmol photons m^-2^ s^-1^) in the presence of 15 µM 3-(3,4-dichlorophenyl) 1,1-dimethylurea (DCMU).

Two new sub-samples (measured independently) were then exposed for 2 min to a 40 µmol photons m^-2^ s^-1^ actinic light (red light, 620 nm) to activate photosynthesis and then, for the next 3 min, one sub-sample remained under 40 µmol photons m^-2^ s^-1^ (control) while the other was exposed to a high light (HL) of 570 µmol photons m^-2^ s^-1^. After 3 min, a pseudo-steady-state was reached and a series of measurements were performed sequentially as fast as possible (8 min in total, with at least 30 s between measurements that require a saturating pulse). The actinic illumination was kept constant except during short dark windows (<500 µs) required along the various measurements. Steady-state (*F_s_*) and maximum PSII fluorescence (*F_M_*’) under light were measured with a detecting light pulse before and after a 200 ms saturating pulse, respectively. The quantum yields of the three energy dissipation pathways in PSII were calculated as: PSII photochemical conversion (ϕPSII) = (*F_M_*’-*F_s_*)/*F_M_*’, ϕNPQ = *F_s_*/*F_M_*’-*F_s_*/*F_M_* and non-regulated heat losses (ϕNO) = *F_S_*/*F_M_*. The fraction of open reaction centres (qL) were calculated as qL = (*F_M_*’-*F_s_*)/(*F_M_*’-*F_0_*’) * *F_0_*’/*F_s,_* where *F_0_*’ is the minimum fluorescence in a sample during or after light exposure with all Q_A_ oxidized. In a model where NPQ is the result of a Stern-Volmer-like quenching process (NPQ/qZ), *F_0_*’ can be theoretically extrapolated under light as *F_0_*/(*F_M_*-*F_0_*_)_/*F_M_*+*F_0_*/*F_M_*’) and qL thus estimates the redox state of Q_A_ (i.e. the concentration of photochemical quencher) (Kramer et al., 2004). A shorter pulse sequence (22 ms) was used to measure P700 redox state at under different conditions: at steady-state under light before the pulse (*P_S_*), at its most oxidized at the end of the pulse (*P_M_*’), and fully re-reduced in the dark after the pulse (*P_0_*). The yield of PSI photochemistry was calculated as ϕPSI = (*P_S_*-*P_M_*’)/(*P_o_*-*P_M_*’), and PSI donor side limitation (ϕND) as = -*P_0_*/*P_M_* and PSI acceptor side limitation (ϕNA) as = 1-ϕPSI-ϕND. Relative electron flow at PSII (rETR_II_= ϕPSII\**E*) estimates the linear electron flux (LEF) while relative electron flow at PSI (rETR_I_= ϕPSI\**E*) is the sum of LEF and cyclic electron flow (CEF), or total electron flow (TEF). There was no effect of Lhcx1 accumulation on PS antenna functional size among the strains (Supplementary Fig. S5) and in diatoms there are no rapid changes in absorption cross-sections (i.e., state-transition) (Lepetit et al., 2022). Therefore, relative changes in the contribution of CEF to LEF due to NPQ can be estimated by comparing the yields of both PS in condition with and without NPQ (570 and 40 µmol photons m^-2^ s^-1^, respectively) as:

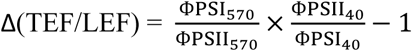

For example, a Δ(TEF/LEF) of 0.4 in steady state under 570 µmol photons m^-2^ s^-1^ indicates that CEF contribution to TEF has increased by 40%, from the unknown — but roughly equal among strains — baseline CEF contribution to TEF at 40 µmol photons m^-2^ s^-1^. The same rationale can be used to compare CEF and LEF at the same light intensities but without and with NPQ, e.g. during its relaxation (see below). In a third sequence, the actinic light was shut off (during 175 ms, averaged 7 times with 2 s dead time between curves) to measure the photochemical rate (turnover rate of both photosystems) via the ECS_lin_ decay. At the light shut-off only PSI and PSII stop contributing to the electric potential gradient instantaneously, therefore the initial slope in ECS_lin_ decay reflect the photochemical rate per PS (Bailleul et al., 2010a). The ECS_lin_ decay was fitted as ECS_T_\**e*^(-*t*/^*^τ^*^)^ where ECS_T_ is an approximation of the light induced proton motive force (Avenson *et al.,* 2005), τ is the lifetime and the initial slope is calculated as ECS_T_/τ. The number of oxidized *c*-t cyt. under 570 µmol photons m^-2^ s^-1^ was estimated as the amplitude of the change in the *c*-t cyt. re-reduction signal during 175 ms dark window, normalized to the laser flash signal. This whole sequence of measurements was conducted on the 10 strains, and on three independent biological samples for each.

For three strains, WT, Lhcx1-Ko and one Lhcx1-OE (LtpW), a similar experiment was conducted but for which 10 sub-samples per strain were exposed to different light intensities (9 steps from 0 to 1000 µmol photons m^-2^ s^-1^). In this case, an additional saturating pulse sequence was used to estimate the maximum proportion of *c*-t cyt. which can be oxidized with a saturating pulse superimposed to the actinic illumination, allowing to normalized *c*-t cyt.^+^ to the fraction of total *c*-t cyt. This experiment served as rationale to identify 570 µmol photons m^-2^ s^-1^ as the optimal light intensity to maximize NPQ contrasts between strains.

For the *transient NPQ relaxation and CEF experiment*, four strains (WT, Lhcx1-Ko and two Lhcx1-OE (LtpW and LtpM)) were prepared as described above before measurements. In samples for which NPQ/qZ was gradually induced to its maximum value, the actinic light was abruptly switched down to 26 µmol photons m^-2^ s^-1^. Under this lower, light-limiting intensity, the relaxation of qZ and the recovery of ϕPSII was monitored in one sub-sample. The experiment was repeated to measure ϕPSI during the relaxation of qZ in another sub-sample from the same biological replicate. The relaxation kinetics of Δ(TEF/LEF) could be calculated as described above for all time points and replacing ϕPSI_40_ by the values measured when qZ had fully relaxed.

### Photoinhibition and NPQ deconvolution analysis

To determine if the rapid and reversible qZ component of NPQ protects PSII against photoinhibition, a prolonged HL stress experiment was conducted in a fluorescence imaging system equipped with a green (532 nm) actinic light (SpeedZen, JbeamBio, France), allowing to measure multiple 60 µL samples in parallel on a same plate. Six strains, two low, two medium and two high Lhcx1 expressing strains, including WT and Lhcx1-Ko, were selected and prepared as described above. The samples were first exposed to 4 min of low light (35 µmol photons m^-2^ s^-1^ Speedzen setting, giving ≈ 15 μmol photons m^-2^ s^-1^ measured at the plate level with a US-SQS/L spherical quantum sensor (Walz, Germany)) and then let to relax in the dark 1 min before the maximum PSII yield in the dark was measured (*F_V_*/*F_M_*) (Fig. S6). Samples were then illuminated under HL (750 µmol photons m^-2^ s^-1^ Speedzen setting, giving ≈ 600 µmol photons m^-1^ s^-1^ measured at the plate level) for 60 min and PSII fluorescence parameters were measured every 3 min. Samples were then let to recover over six low light/dark intervals (4: 1 min) identical to before the stress, during which PSII fluorescence parameters were measured every min. This protocol accelerates qZ relaxation and promotes PSII repair under low light, and allow to compare *F_V_*/*F_M_* before light stress and at its maximal recovered value (*F_V_*’/*F_M_*’Rec.) in standardized conditions that should allow to fully oxidized Q_A_ after 1 min of darkness. The illumination protocol is summarized in (Supplementary Fig. S7). For each repetition of this experiment, each biological replicate was measured in six sub-samples in parallel, supplemented with increasing final concentration (0 to 0.8 mg mL^-1^, 5 min before the start of the experiment) of lincomycin (Sigma-Aldrich), which inhibits chloroplast translation and therefore *de novo* synthesis of PsbA for PSII repair. The average of treatments at 0.4 and 0.8 mg mL^-1^ lincomycin (saturated effect on *F_V_*’/*F_M_*’Rec. (see Supplementary Fig. S8)) were used as comparison with the control conditions.

Under moderate light stress, *P. tricornutum* exhibits a single qZ component of NPQ, proportional to DT bound to Lhcx1 and which relax under ≈10 min under low light or darkness (Croteau et al., 2025). Under longer light stress, photodamaged PSII centres cannot participate to electron transport but can act as nonphotochemical quenchers (qI), complexifying the relationship between total NPQ and *F_V_*’/*F_M_*’ (Nawrocki et al., 2021). To clarify the role of qZ in protecting PSII from photoinhibition and qI in diatoms, NPQ relaxation was fitted as the sum of a fast (qZ) and a slow (qI) monoexponential decay.

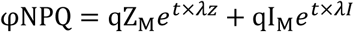

Where qZ_M_ and qI_M_ are the maximal contribution of each component to ϕNPQ, i.e., at the beginning of the relaxation, and λ_Z_ and λ_I_ are the decay rates of each component (see Supplementary Fig. S9).

### High light experiment for biochemistry and RNA sampling

A 60 min HL (600 µmol photons m^-2^ s^-1^) stress followed by 30 min of recovery under low light (15 µmol photons m^-2^ s^-1^) experiment was reproduced in an experimental setup allowing to work on large volumes of non-concentrated cultures always at 1*10^6^ c/mL (Erlen maintained agitated over a LED panel) and to sample biological material. To monitor PSII fluorescence parameters in parallel of these experiments, 1 mL sub-samples were let to relax 1 min in the dark in a home-made fluorometer before measing minimum and maximal fluorescence, *F_0_*’ and *F_M_*’, respectively. The reversible-qZ portion of NPQ was estimated from the highest recovered value of *F_M_*’ post-stress as reference *F_M_*’Rec., with qZ = *F_M_*’Rec./*F_M_*’-1. The maximal qZ measured in given replicate over the course of an experiment is used to define the colour scale in Fig. 2 to Fig. 4, and as *x*-axis coordinate of each time-panel for the gene expression profiles. To examine the relationship between PsbA degradation, qI, and *F_V_*’/*F_M_*’Rec., three strains (WT, Lhcx1-Ko, and two Lhcx1-OE lines) were grown and maintained in exponential phase in 100 mL cultures. A 10 mL protein sample was collected at the initial time point (*t_0_*). The remaining culture was then split into two aliquots: one supplemented with lincomycin (final concentration 0.4 mg mL⁻¹) and one left untreated. For both conditions, an additional 10 mL sample was collected at the end of the experiment (after 30 min recovery under low light (15 µmol photons m^-2^ s^-1^). Samples were transferred to 15 mL Falcon tubes, centrifuged for 10 min at 2,500 rpm at 4 °C, rinsed in PBS, and stored at −80 °C.

For RNA sampling, the same experiment (without lincomycin) was repeated using 350 mL cultures of six strains. At *t_0_*, *t_20_*, and *t_60_* of the HL treatment, 100 mL aliquots were filtered onto Whatman 589/2 filters and immediately froze in liquid nitrogen. At the same time points, 10 mL samples for pigment analysis were collected, centrifuged, and stored at −80 °C.

### Immunoblotting

Immunodetection of PsbA was performed following the same protocol as in (Giovagnetti et al., 2022), using rabbit-polyclonal PsbA (AS05 084, 1: 50,000) and ATPsynthase-βCF_1_ (AS03 030, 1: 10,000 (loading control)) antibodies (Agrisera, Sweden). Total protein loading was of 10 µg to ensure the detection of both antibodies was in the linear range.

### RNA extraction and analysis of transcriptomic data

Three biological replicates of six different strains with ranging maximal NPQ/qZ values were selected for the HL transcriptomic experiment. Three different timepoints (0, 20 and 60 min) were sampled for transcriptomic analysis. Total RNA was extracted with TriPure Isolation Reagent (Roche) following the protocol provided by the company, further purified by an ammonium acetate precipitation, and finally treated with RQI DNaseI (Promega). Libraries of cDNA were prepared from extracts enriched in mRNA by oligo-T selection and sequenced by the DNBseq platform at the Beijing Genomics Institute (*BGI* Group). Sequencing, mapping over *P. tricornutum* second version genome (Phatr, and transcripts quantification of the extracted RNA samples was outsourced to BGI (Hong Kong). We retrieve 4.35Gb bases per sample with an average mapping using HISAT2 of 90.26 % on PHATR2 genome (74 % uniquely mapped) (Supplementary Table S1). A total of 50 transcriptomes was retrieved because one replicate of LtpW started to revert during culturing and one sample of Lhcx1-Ko *t*_0_ was lost during RNA extraction. Among the 10,381 genes identified over RNA sequencing (BGI), 8,710 were kept for further analysis, as genes with at least two samples showing very low FPKM (<10) were removed. The large number of transcriptomes analysed allowed to develop a novel approach to decipher genes whose expression respond directly to *absolute* light intensity (along a light exposure time dimension) from those responding to photosynthetic signals deeply influenced by NPQ/qZ/, i.e. *experienced* light (shown in Fig. 4).

For every transcript, mRNA abundance was quantified and first normalized by fragments per kilobase of transcript per million fragments mapped (FPKM) for each sample at *t*_0_, *t*_20_ and *t*_60_ min. A min-max normalization was then applied to FPKM values across all 50 transcriptomes so that the values of each transcript are bounded between 0 and 1, these rescaled values are referred to as *normed FPKM* in Fig. 4a. The average expression of the 16-18 samples at each time point represents the time dependence of the HL-induced changes. The standard deviation (σ) of these three averaged values at *t*_0_, *t*_20_ and *t*_60_ was calculated and indicated as σ_HL_ in Fig. 4a. The NPQ/qZ effect is calculated by subtracting time dependence of the HL-induced changes to the *normed FPKM* at each time point. The standard deviation of the 50 values after this subtraction was calculated and indicated as σ_NPQ/qZ_ in Fig. 4a. With the objective that the clustering method gives equal importance to time and NPQ/qZ dependences, we modified gene expression profiles by assigning the same weight to both dependencies. For this, the time dependence and the NPQ/qZ dependence were divided by the average value of σ_HL_ andσ_NPQ/qZ_, respectively. Finally *weighted FPKM* are obtained by combining both normalized time and NPQ/qZ dependencies.

For K-means clustering, random seeding was used. Data was organized into three discrete HL exposure snapshots: 0, 20 and 60 min, ordered from left to right, with samples ranked from lowest to highest NPQ/qZ capacity — as empirically determined during the experiment — within each time panel. A Silhouette analysis supported six as the optimal number of clusters. Gene Ontology (GO) enrichment analyses were performed using the *compareCluster*() function from the clusterProfiler package in R, with a P-value cutoff of 0.05. Only the enriched GO categories are shown in Fig. 4 and Supplementary Fig. S12. The GO enrichment network displayed in Supplementary Fig. S13 was generated using the *emaplot*() function from the enrichplot package.

## Supporting information

Croteau_et_al_Supp

## Acknowledgements

The authors want to thank Pascal Campagne and Bernard Lepetit for fruitful discussions about the transcriptomic data and bioinformatic analysis. This research was supported by the European Research Council (ERC) PhotoPHYTOMIX project (grant agreement No. 715579) and the ANR ‘BrownCut’ (ANR-19-CE20-0020). AF was supported by funding from the Fondation Bettencourt-Schueller (Coups d’élan pour la recherche francaise-2018) and the “Initiative d’Excellence” program (Grant “DYNAMO,” ANR-11-LABX-0011-01).

## Abbreviations

AEF: Alternative Electron Flow
ATP: Adenosine Triphosphate
CEF: Cyclic Electron Flow
*c*-t cyt.: c-type cytochromes (cytchrome *c_6_* + cytochrome *f*)
DD: Diadinoxantin
DT: Diatoxanthin
DNA: Deoxyribonucleic acid
ECS: Electrochromic Shift
FCP: fucoxanthin chlorophyll-a/c protein
FPKM: fragments per kilobase of transcript per million fragments mapped
*F_V_*/*F_M_*: Maximum yield of PSII photochemistry in the dark
*F_V_^’^/F_M_^’^*: Maximum yield of PSII photochemistry in the dark after light exposure
GO: Gene Ontology
HL: High ligh
LEF: Linear Electron Flow
Lhcf: Light-Harvesting Complex F
Lhcr: Light-Harvesting Complex R
Lhcx: Light-Harvesting Complex X
NPQ: Non-Photochemical Quenching of chlorophyll fluorescence
P700: Photosystem I primary donor
PCA: Principal Component Analysis
PQ/PQH_2_: Plastoquinone/Plastoquinol
PsbA/PsbB: Core proteins forming PSII heterodimer
PSI: Photosystem I
PSII: Photosystem II
Pt: *Phaeodactylum tricornutum*
ROS: Reactive Oxygen Species
qI: Photoinhibition-related quenching
qL: Fraction of open PSII reaction centres
qT: State transitions-dependent quenching
qZ: Xanthophyll-dependent quenching
TEF: Total Electron Flow
^t^RNA: Transfer ribonucleic acid
VDE: Violaxanthin deepoxidase
WT: wildtype
XC: Xanthophyll cycle
ZEP3: Zeaxanthin epoxidase 3
Δ(TEF/LEF): Relative change in total electron flow proportion versus linear electron flow
ΦPSII: Yield of PSII photochemistry,
ΦNPQ: Yield for dissipation by NPQ downregulation
ΦNO: Yield of other non-photochemical losses
ΦPSI: Yield of PSI photochemistry
ΦND: PSI yield of non-photochemical energy dissipation due to donor side limitation
ΦNA: PSI yield of non-photochemical energy dissipation due to acceptor side limitation

## Notes

### Competing Interest Statement

The authors have declared no competing interest.

