## Supplementary material for "A photoprotection dial maps holistic light-stress response in diatoms": Croteau_et_al_Supp

### Supplementary materials

Supplementary Table S1

Supplementary Figure S1-S13

**Supplementary Table S1| Table grouping the strains used for the transcriptomic experiment.** Listed for each strain are the relative Lhcx1 abundance, the empirically measured maximal NPQ/qZ reached during the experiment (per biological replicate), and key read-mapping quality metrics.

| Strains | Rep | Sample | Tot_Clean | MapRatio | Uniquely.MappingRatio | Strand.Specific.Ratio | t | Lhcx1 (r.u.) | Maximum NPQ/qZ |
|---|---|---|---|---|---|---|---|---|---|
| Ko | 1 | Ko_t0_1 | 43312162 | 0.910 | 0.732 | 0.924 | t0 | 0.033 | 0.373 |
| Ko | 1 | Ko_t20_1 | 44006896 | 0.916 | 0.757 | 0.937 | t20 | 0.033 | 0.373 |
| Ko | 1 | Ko_t60_1 | 42279670 | 0.914 | 0.753 | 0.927 | t60 | 0.033 | 0.373 |
| Ko | 2 | Ko_t20_2 | 43915104 | 0.913 | 0.743 | 0.927 | t20 | 0.033 | 0.228 |
| Ko | 2 | Ko_t60_2 | 43903864 | 0.911 | 0.746 | 0.933 | t60 | 0.033 | 0.228 |
| Ko | 3 | Ko_t0_3 | 43461984 | 0.904 | 0.725 | 0.914 | t0 | 0.033 | 0.414 |
| Ko | 3 | Ko_t20_3 | 43827130 | 0.913 | 0.755 | 0.915 | t20 | 0.033 | 0.414 |
| Ko | 3 | Ko_t60_3 | 44128682 | 0.910 | 0.756 | 0.907 | t60 | 0.033 | 0.414 |
| WT | 1 | WT_t0_1 | 42991984 | 0.919 | 0.741 | 0.924 | t0 | 0.180 | 1.339 |
| WT | 1 | WT_t20_1 | 44146500 | 0.919 | 0.750 | 0.944 | t20 | 0.180 | 1.339 |
| WT | 1 | WT_t60_1 | 43996630 | 0.908 | 0.739 | 0.912 | t60 | 0.180 | 1.339 |
| WT | 2 | WT_t0_2 | 42545516 | 0.917 | 0.741 | 0.895 | t0 | 0.180 | 1.261 |
| WT | 2 | WT_t20_2 | 44137450 | 0.918 | 0.753 | 0.913 | t20 | 0.180 | 1.261 |
| WT | 2 | WT_t60_2 | 44052582 | 0.912 | 0.746 | 0.916 | t60 | 0.180 | 1.261 |
| WT | 3 | WT_t0_3 | 43204272 | 0.920 | 0.751 | 0.909 | t0 | 0.180 | 1.426 |
| WT | 3 | WT_t20_3 | 44037818 | 0.915 | 0.761 | 0.910 | t20 | 0.180 | 1.426 |
| WT | 3 | WT_t60_3 | 44265028 | 0.907 | 0.759 | 0.928 | t60 | 0.180 | 1.426 |
| LtpN | 1 | N_t0_1 | 43273034 | 0.902 | 0.726 | 0.914 | t0 | 0.233 | 1.824 |
| LtpN | 1 | N_t20_1 | 44062924 | 0.904 | 0.745 | 0.936 | t20 | 0.233 | 1.824 |
| LtpN | 1 | N_t60_1 | 42215848 | 0.905 | 0.746 | 0.942 | t60 | 0.233 | 1.824 |
| LtpN | 2 | N_t0_2 | 43383722 | 0.907 | 0.727 | 0.909 | t0 | 0.233 | 1.155 |
| LtpN | 2 | N_t20_2 | 43808750 | 0.905 | 0.740 | 0.916 | t20 | 0.233 | 1.155 |
| LtpN | 2 | N_t60_2 | 44211730 | 0.903 | 0.738 | 0.932 | t60 | 0.233 | 1.155 |
| LtpN | 3 | N_t0_3 | 43106024 | 0.903 | 0.733 | 0.890 | t0 | 0.233 | 1.915 |
| LtpN | 3 | N_t20_3 | 43903808 | 0.910 | 0.753 | 0.913 | t20 | 0.233 | 1.915 |
| LtpN | 3 | N_t60_3 | 44087164 | 0.905 | 0.749 | 0.932 | t60 | 0.233 | 1.915 |
| LtpY | 1 | Y_t0_1 | 43281490 | 0.907 | 0.726 | 0.915 | t0 | 0.616 | 2.698 |
| LtpY | 1 | Y_t20_1 | 43988322 | 0.903 | 0.733 | 0.907 | t20 | 0.616 | 2.698 |
| LtpY | 1 | Y_t60_1 | 43574182 | 0.896 | 0.733 | 0.907 | t60 | 0.616 | 2.698 |
| LtpY | 2 | Y_t0_2 | 43275170 | 0.903 | 0.725 | 0.894 | t0 | 0.616 | 2.473 |
| LtpY | 2 | Y_t20_2 | 44096796 | 0.909 | 0.746 | 0.912 | t20 | 0.616 | 2.473 |
| LtpY | 2 | Y_t60_2 | 43948298 | 0.896 | 0.734 | 0.914 | t60 | 0.616 | 2.473 |
| LtpY | 3 | Y_t0_3 | 43276056 | 0.906 | 0.737 | 0.871 | t0 | 0.616 | 2.318 |
| LtpY | 3 | Y_t20_3 | 44259002 | 0.903 | 0.757 | 0.897 | t20 | 0.616 | 2.318 |
| LtpY | 3 | Y_t60_3 | 44218736 | 0.897 | 0.754 | 0.919 | t60 | 0.616 | 2.318 |
| LtpW | 1 | W_t0_1 | 43230568 | 0.900 | 0.712 | 0.923 | t0 | 0.862 | 2.125 |
| LtpW | 1 | W_t20_1 | 43803456 | 0.890 | 0.715 | 0.917 | t20 | 0.862 | 2.125 |
| LtpW | 1 | W_t60_1 | 43885252 | 0.881 | 0.713 | 0.925 | t60 | 0.862 | 2.125 |
| LtpW | 2 | W_t0_2 | 43325016 | 0.901 | 0.718 | 0.907 | t0 | 0.862 | 3.973 |
| LtpW | 2 | W_t20_2 | 44007742 | 0.894 | 0.729 | 0.916 | t20 | 0.862 | 3.973 |
| LtpW | 2 | W_t60_2 | 42301782 | 0.886 | 0.721 | 0.927 | t60 | 0.862 | 3.973 |
| LtpM | 1 | M_t0_1 | 43343822 | 0.901 | 0.723 | 0.910 | t0 | 0.934 | 3.613 |
| LtpM | 1 | M_t20_1 | 44250858 | 0.895 | 0.734 | 0.935 | t20 | 0.934 | 3.613 |
| LtpM | 1 | M_t60_1 | 42239214 | 0.885 | 0.729 | 0.940 | t60 | 0.934 | 3.613 |
| LtpM | 2 | M_t0_2 | 42826340 | 0.901 | 0.721 | 0.898 | t0 | 0.934 | 3.936 |
| LtpM | 2 | M_t20_2 | 43514828 | 0.898 | 0.731 | 0.916 | t20 | 0.934 | 3.936 |
| LtpM | 2 | M_t60_2 | 43760758 | 0.884 | 0.717 | 0.906 | t60 | 0.934 | 3.936 |
| LtpM | 3 | M_t0_3 | 42766332 | 0.899 | 0.729 | 0.890 | t0 | 0.934 | 3.646 |
| LtpM | 3 | M_t20_3 | 43832942 | 0.895 | 0.737 | 0.901 | t20 | 0.934 | 3.646 |
| LtpM | 3 | M_t60_3 | 44017100 | 0.883 | 0.730 | 0.924 | t60 | 0.934 | 3.646 |

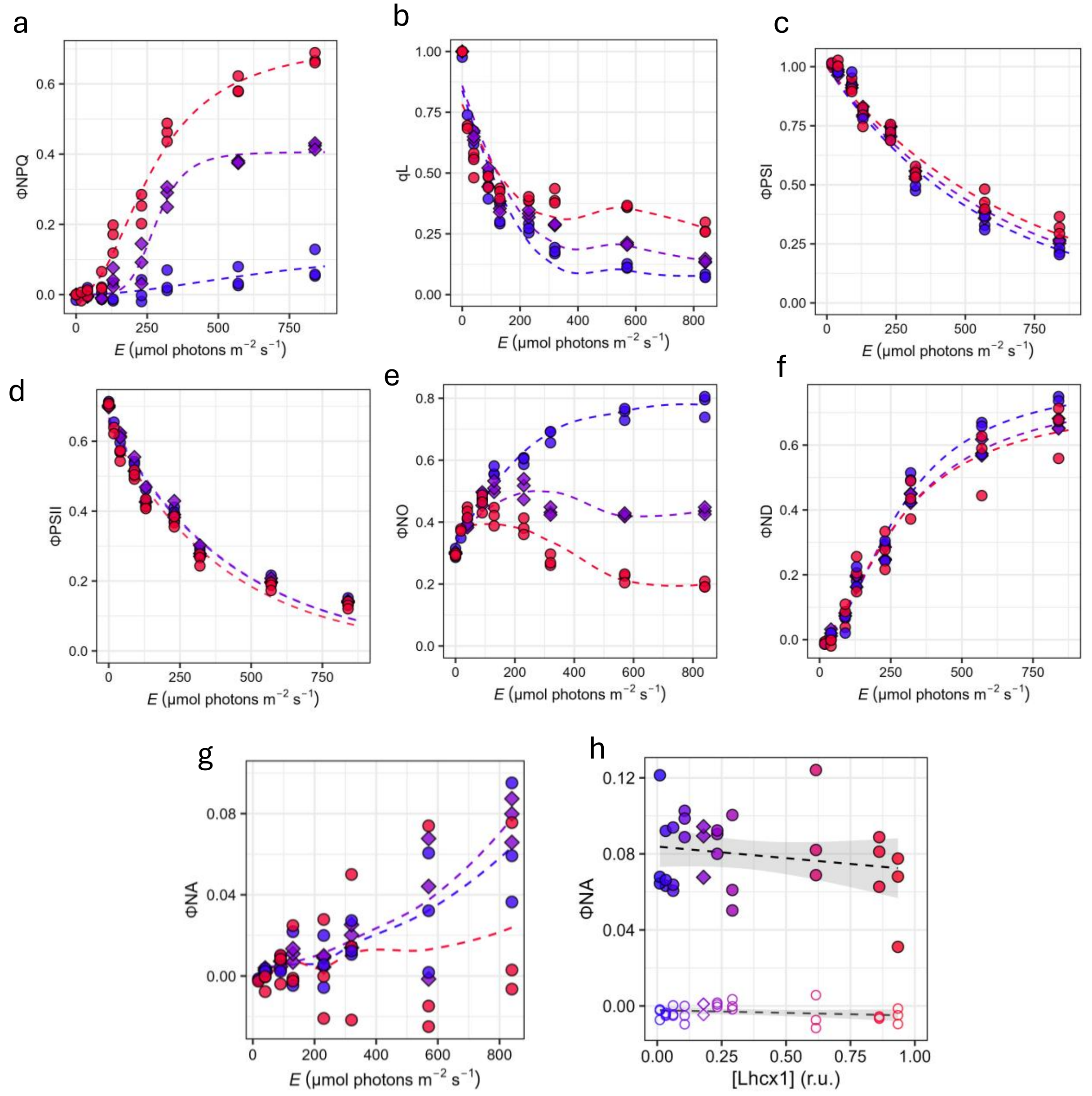


**Supplementary Fig. S1| NPQ induction modulates the entire electron transport chain bioenergetics in *Phaeodactylum tricornutum*.** This figure reports the same parameters as presented in Fig. 1 of the main manuscript but as a function of light intensity (*E*), in three strains, at contrast of Fig. 1 which examined 10 strains under two light intensities. The strains used are Pt2-WT (diamond symbols), the Lhcx1-Ko, and an overexpressor LtpW, with Lhcx1 abundance (correlated to NPQ capacity) color-couded from blue to red. **Top row**: the quantum yield of NPQ (ϕNPQ) (a), qL (which approximates $Q_A$ redox state in the lake model, and thus provides a proxy for the plastoquinone pool ($PQ/PQH_2$) redox state) (b) and the quantum yield of photosystem I (ϕPSI) (c). **Bottom row**: ϕPSII (d), yield of non-regulated heat dissipation by PSII (ϕNO) (e) and the yield of PSI donor-side limitation (ϕND, i.e. the proportion of oxidized PSI-P700) (f). Additionally, the yield of PSI acceptor-side limitation, which we lack space to show in Fig. 1, is shown as a function of *E* (g) and as a function of Lhcx1 concentration across the 10 strains (open symbols: 40 µmol photons $m^{-2}$ $s^{-1}$; closed symbols: 570 µmol photons $m^{-2}$ $s^{-1}$) (h).

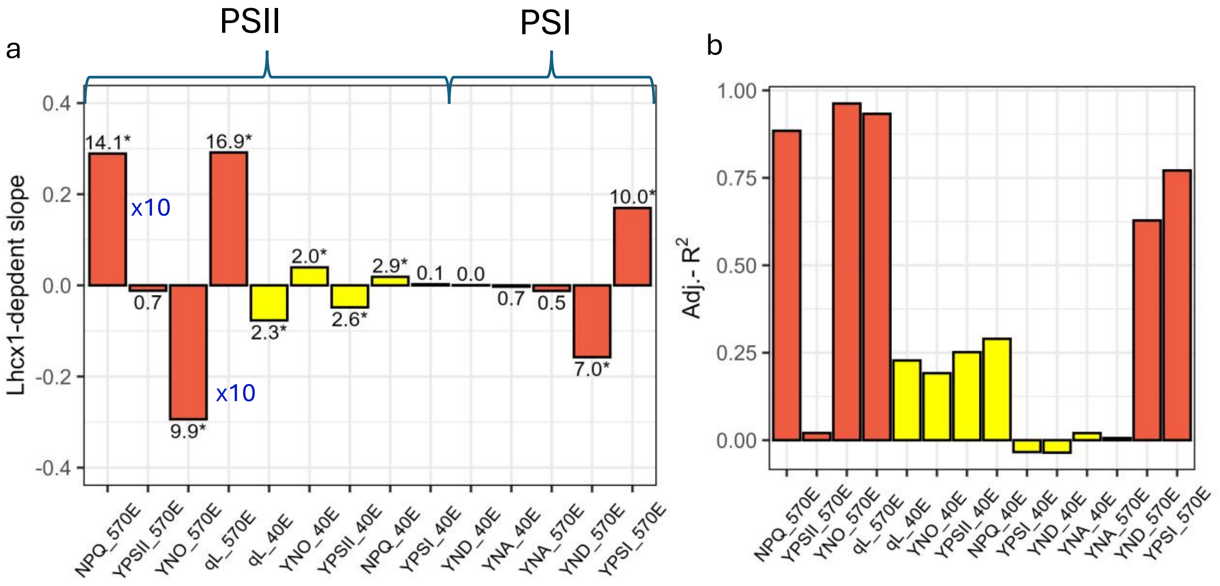


**Supplementary Fig. S2 | Statistical summaries of parameters shown in Fig. 1 fitted as a function of Lhcx1 concentration.** Parameters are fitted under high light (HL; 570 µmol photons $m^{-2}$ $s^{-1}$, red bars) and low light (LL; 40 µmol photons $m^{-2}$ $s^{-1}$, yellow bars) illumination (all parameters' plot are show in either Fig. 1 or Supplementary Fig. S1). Linear slope coefficients are shown for all parameters, except for the quantum yield of PSII non-regulated heat dissipation (YNO) under HL, which was fitted with a monoexponential decay; its initial rate of decrease (linearized at Lhcx1 = 0) is shown here. For clarity, the HL slopes for NPQ and YNO under HL are divided by 10. The $\log_{10}$(*P*-value) is indicated above each bar, with * denoting $P < 0.05$. Adjusted $R^2$ values for each relationship are shown in (b) .

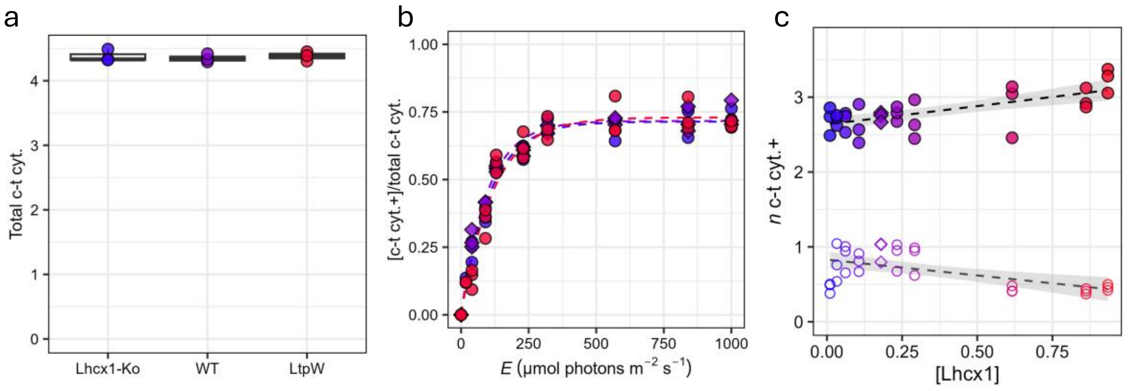


**Fig. S3 | Light-induced oxidation of c-type cytochromes (c-t cyt.) and its modulation by Lhcx1-dependent qZ.** Based-off absorbance change measurements with the JTS normalized to signal amplitude corresponding to a single charge separation per PSI (a) shows the maximum number of *c*-t cyt. that can be oxidized in the dark by a saturating pulse. (b) Proportion of oxidized *c*-t cyt. under steady-state illumination as a function of light intensity (*E*) in Lhcx1-KO, WT (diamond symbols), and the Lhcx1 overexpressor (LtpW), showing that *c*-t cyt. oxidation saturates at 570 µmol photons $m^{-2}$ $s^{-1}$. (c) Number of *c*-t cyt.+ per PSI across the 10 strains with varying Lhcx1 concentrations, measured under two light intensities (open symbols: 40 µmol photons $m^{-2}$ $s^{-1}$; closed symbols: 570 µmol photons $m^{-2}$ $s^{-1}$) All panels show three independent biological replicates, except one strain in panel (c) for which *n* = 2 under lower light.

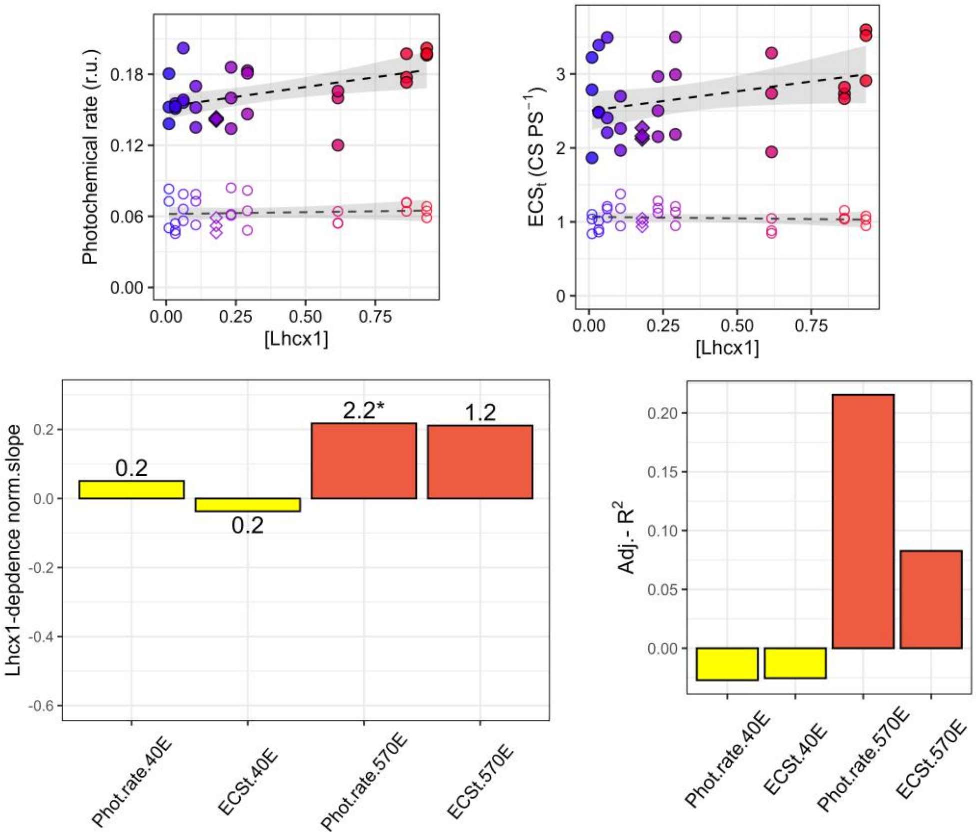


**Supplementary Fig. S4 | Statistical summaries of the photochemical rate and the total ECSt fitted as a function of Lhcx1 concentration.** Parameters are fitted under high light (HL; 570 µmol photons $m^{-2}$ $s^{-1}$, red bars) and low light (LL; 40 µmol photons $m^{-2}$ $s^{-1}$, yellow bars) illumination. In (a) the photochemical rate is the initial slope of linear ECS decay upon light shut-off and in (b) ECSt is total ECS, representing a proxy of the light-induced proton motive force. Linear slope coefficients are shown in (c) and adjusted $R^2$ values for each relationship are shown in (d). With * in (c) denoting $P < 0.05$

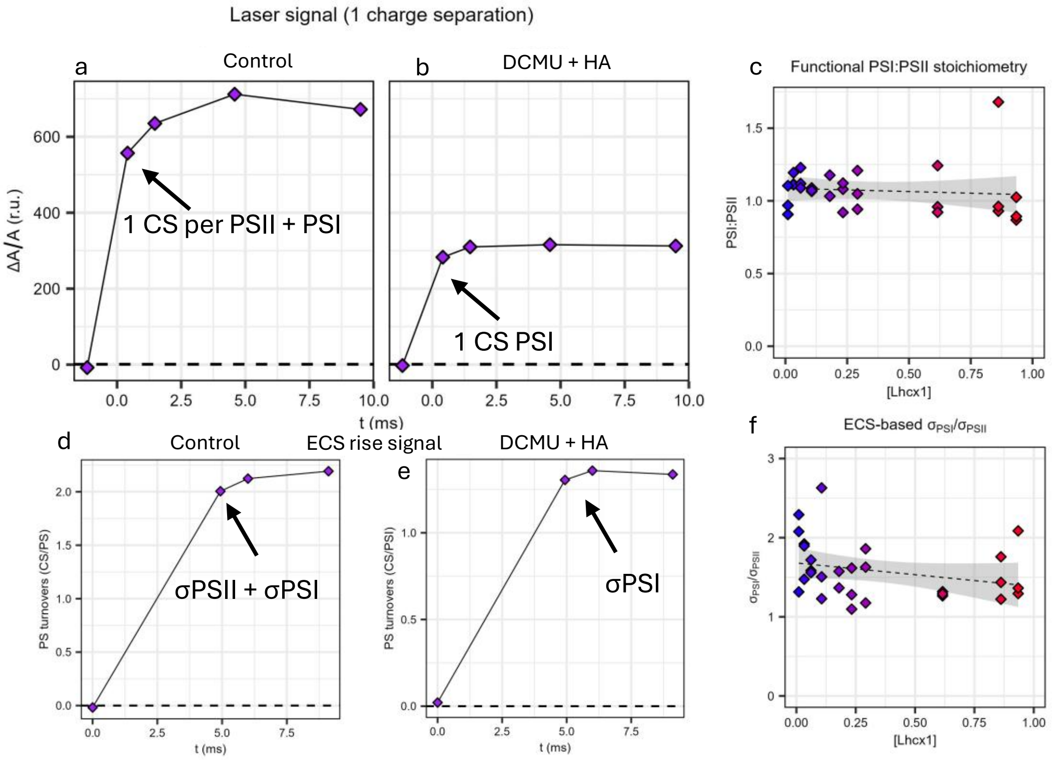


**Supplementary Fig. S5 | Photosystem (PS) I and II stoichiometry and antenna functional cross-section (σ) ratios for ranging Lhcx1 concentration** : Shown only for the WT, the absorbance change signal (ΔA/A) for the linear electrochromic shift ($ECS_{lin}$) following a laser flash inducing a single charge separation (CS) per photosystem (PS), in control conditions (a) and when PSII is inhibited by 3-(3,4-dichlorophenyl) 1,1-dimethylurea (DCMU) and hydroxylamine (HA) (b), and the stoichiometry of functional PSI-to-PSII in the 10 *Phaeodactylum tricornutum* strains with ranging Lhxc1 concentrations used in the paper (c). Shown only for the WT, dark-to-light ECS rise under moderate (130 µmol photons m$^{-2}$ s$^{-1}$) light so that light absorption limits photochemistry and allows to measure the ECS-based functional cross-section (σ) of PSII+PSI (d) and PSI (when PSII is inhibited) in (e). To calculate the ratio between σPSI/σPSII in (f), σPSII was divided by Fv/Fm (0.71±0.01 across strains/replicates) to only take into account the fraction of absorbed light energy by PSII that participates to photochemistry (the rest is lost as heat). Overall, we conclude from these measurements that there is no significant Lhcx1-dependent effects on PSI:PSII stoichiometry or σPSI/σPSII ratios among the 10 strains used. All measurements were repeated on three independent biological replicates (except one strains for which *n* = 2 in (c)).

#### Relative CEF under low-light recovery after HL-stress

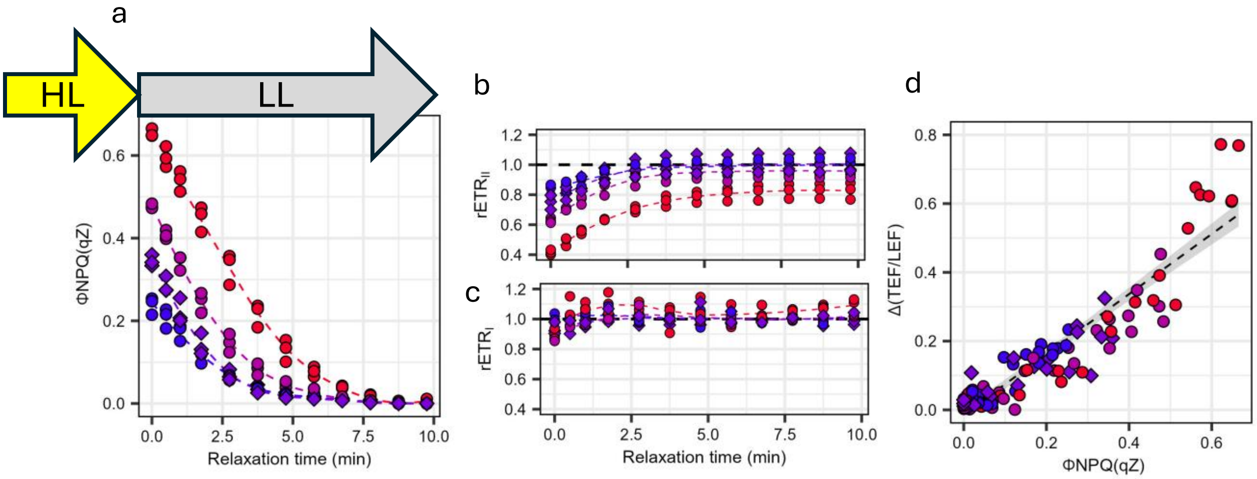


**Supplementary Fig. S6 | Estimation of relative change in cyclic electron flow (CEF) during NPQ/qZ relaxation under low light.** In the main manuscript, we introduced the Δ(TEF/LEF) parameter, which represents the change in total electron flow (TEF; comprising linear electron flow, LEF, and CEF) relative to the change in LEF between conditions with and without NPQ. A positive Δ(TEF/LEF) indicates a relative increase in CEF (see Methods). Because PSII contributes only to LEF, whereas PSI contributes to both LEF and CEF, the relative electron transport rates (rETR) of each photosystem can be used to estimate LEF (from PSII) and TEF (from PSI). Here, the same principle is applied to rETR measured during qZ relaxation under low light (LL; grey arrow), after maximal NPQ/qZ induction under high light (HL; yellow arrow). Four strains were used: Lhcx1-KO, WT (diamond symbols), and two Lhcx1 overexpressors (LtpM and LtpW). (a) Quantum yield of the qZ component of NPQ (ϕNPQ(qZ)), (b) rETRII and (c) rETRI as a function of relaxation time. (d) Resulting Δ(TEF/LEF) values plotted as a function of ϕNPQ(qZ). Three biological replicates were measured for each strain.

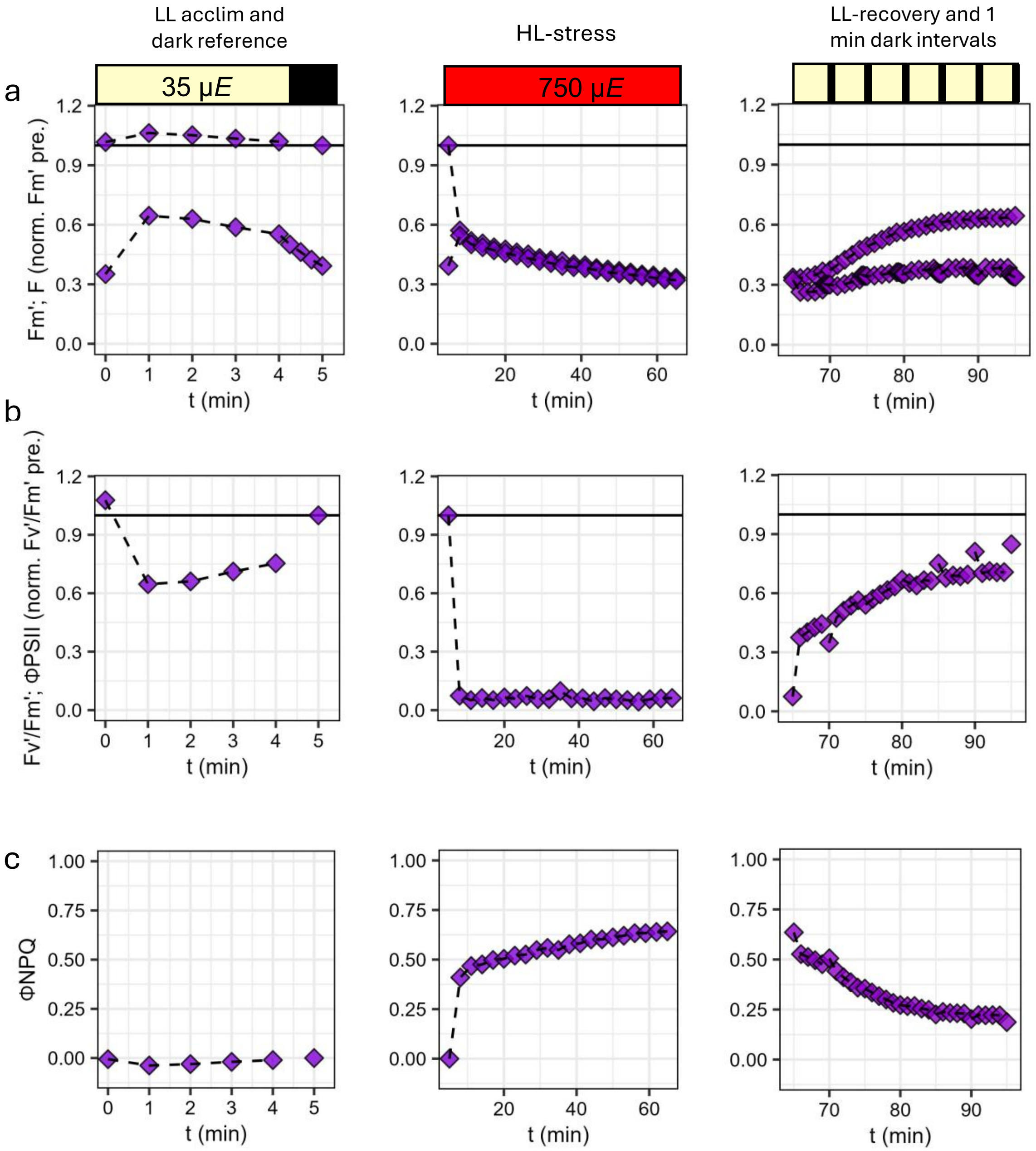


**Supplementary Fig. S7 |Illumination protocol used to estimate the qZ-photoprotection against PSII photoinhibition.** Representation of the illumination protocol (shown for the WT strain without lincomycin) used to test the interaction between non-photochemical quenching (NPQ) and PSII photodamage/photoinhibition. The influence of NPQ-qZ on photoinhibition was assessed from the maximal recovery of the PSII quantum yield in the dark ($F_V'/F_M'$) (Fig. 2f in the main manuscript) and from the amplitude of the slow-relaxing photoinhibitory quenching component ((Fig. 2i in the main manuscript)). **Left column:** Cells were acclimated for 5 min to low light (LL; 35 μmol photons $m^{-2}$ $s^{-1}$), followed by 1 min of darkness to measure the pre-stress fluorescence levels used for normalization after recovery. **Middle column:** Cultures were exposed for 60 min to high light (750 μmol photons $m^{-2}$ $s^{-1}$). **Right column:** Recovery was monitored over 30 min using six alternating LL/dark intervals (4:1), allowing maximal $F_V'/F_M'$ recovery to be measured under conditions comparable to those at the end of LL acclimation. For the three steps of the experiment, i.e. low light acclimation, high light stress and recovery, (a) shows minimum ($F'$) and maximum ($F_M'$) fluorescence, (b) shows $F_V'/F_M'$ or effective quantum yield when expose to light (ϕPSII in the middle column) and (c) shows the quantum yield of NPQ (ϕ). See Methods for details.

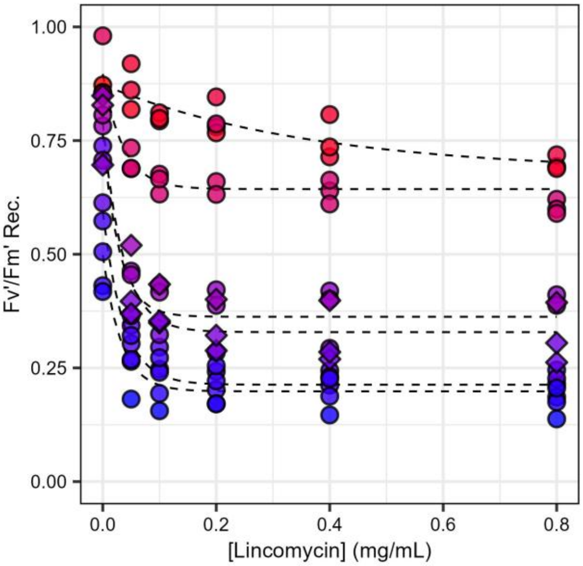


**Supplementary Fig. S8 | Lincomycin dose-effect on high-light induced loss of PSII efficiency**. The quantum yield of PSII in the dark at the end of low light recovery ($F_V'/F_M'$Rec.) following high light stress (see Supplemntary Fig. S7) is shown as a function of the concentration of lincomycin added right before the experiment. From this analysis, we concluded that concentrations 0.4 and 0.8 µg/mL are saturating, and we used these treatments for further analysis shown in Fig. 2. Experimental data correspond to independent biological triplicates in 6 strains (diamond symbols indicate WT).

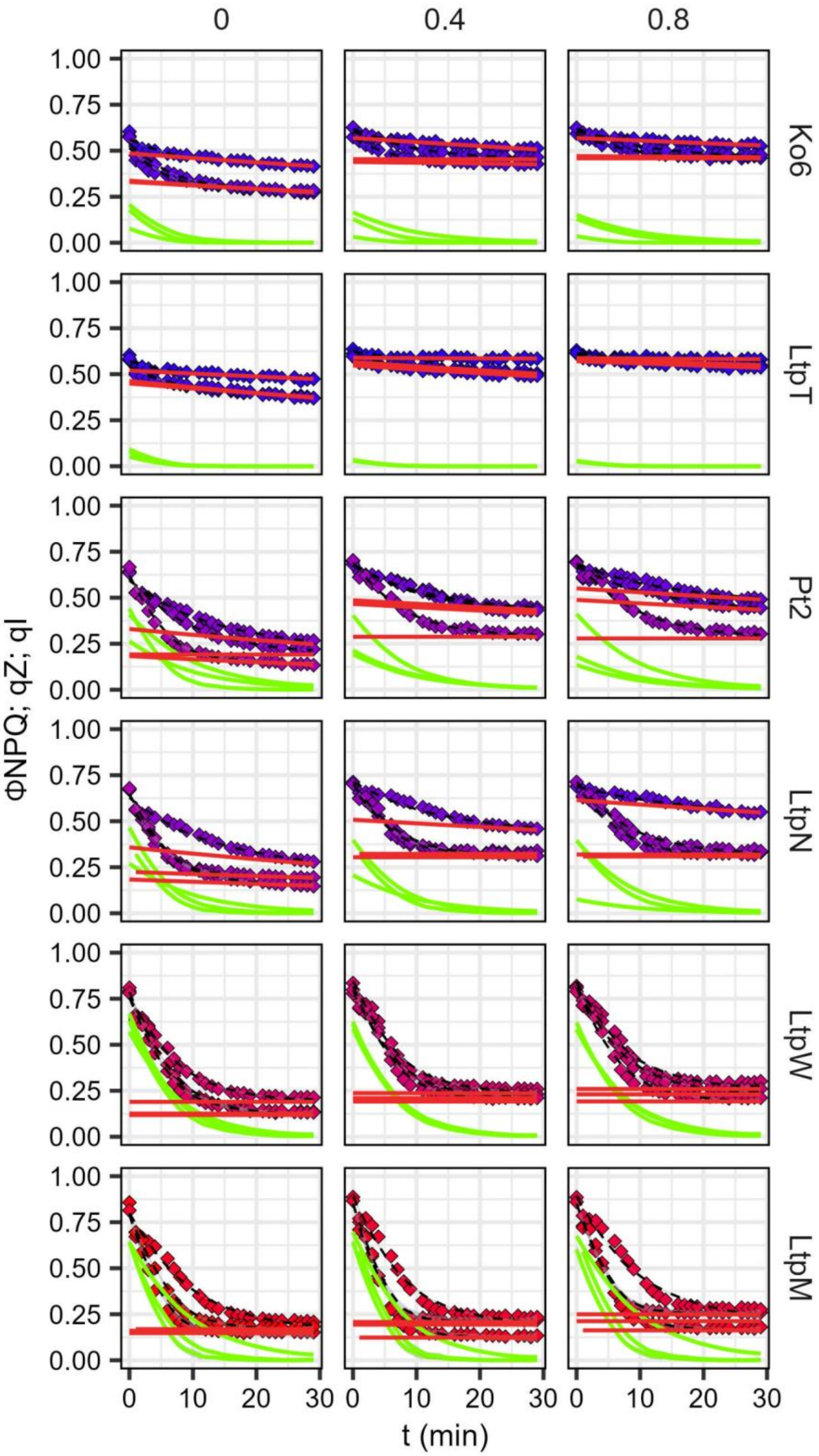


**Supplementary Fig. S9 | Interplay between qZ and qI induced by 1h of high light illumination**. To estimate the respective contributions of Lhcx1-dependent quenching (qZ) and photoinhibition-dependent quenching (qI), the relaxation of ϕNPQ was fitted as the sum of two monoexponential decays, with the fast component attributed to qZ (green line) and the slow component attributed to qI (red line) (see Methods). The fit on the biological triplicates used in each strain are shown for control conditions (left column), and with 0.4 μg mL$^{-1}$ lincomycin (middle column) and 0.8 μg mL$^{-1}$ lincomycin (right column).

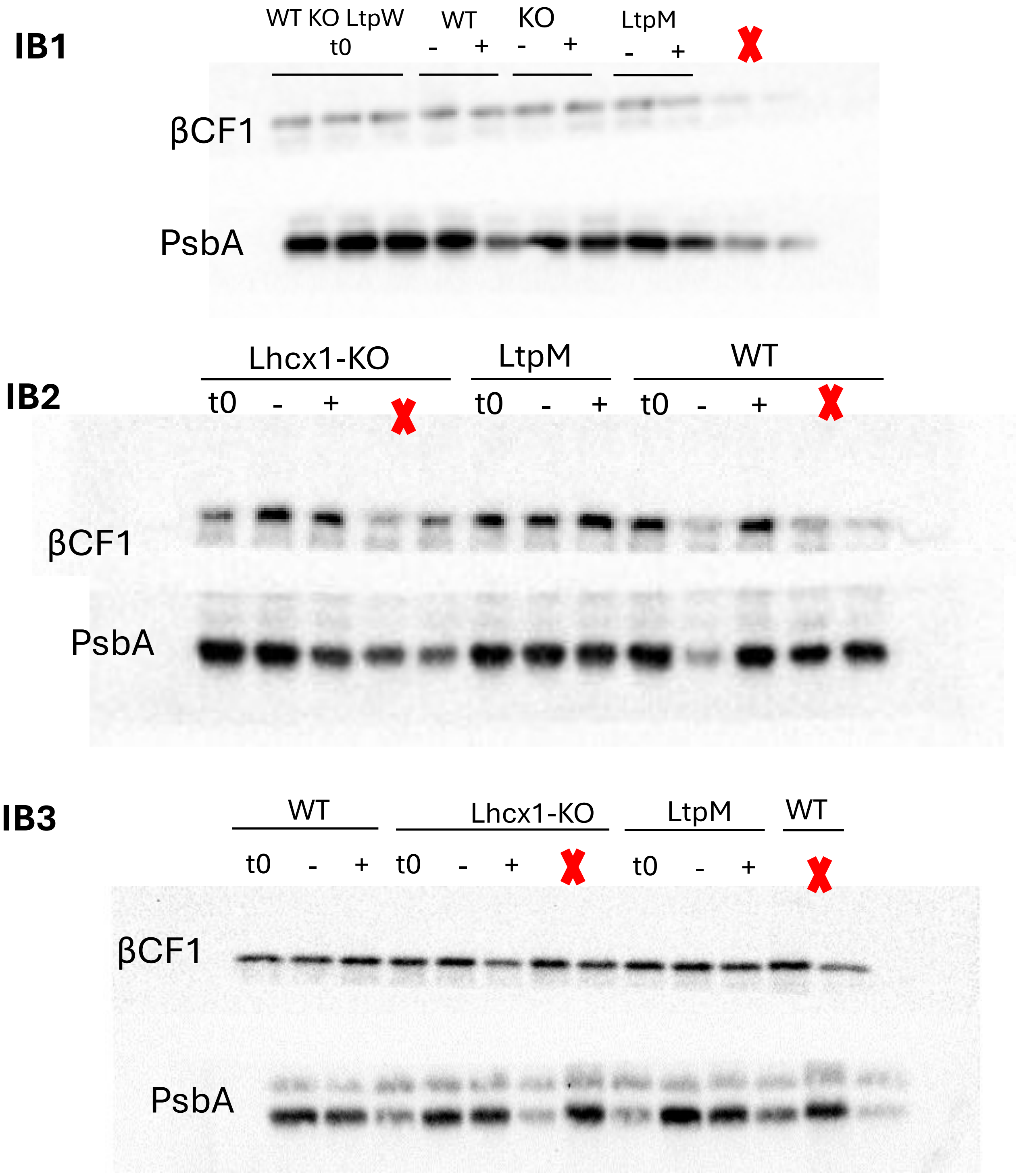


**Supplementary S10| Immunoblots membranes of PsbA quantification before and after high light stress, with and without lincomycin**. Immunoblot (IB) membranes from three independent biological replicates used to quantify the PSII PsbA subunit, with the ATP synthase β$CF_1$ subunit as a loading control. $t_0$ refers to initial conditions before light stress. "+" and "–" indicate samples with and without 0.4 μg $mL^{-1}$ lincomycin, collected after 30 min of low-light recovery following 1 h of high-light stress (see Methods). Red symbols mark extra material that migrated into empty wells but was not used in the analysis. IB3 corresponds to the membrane shown in Fig. 2g.

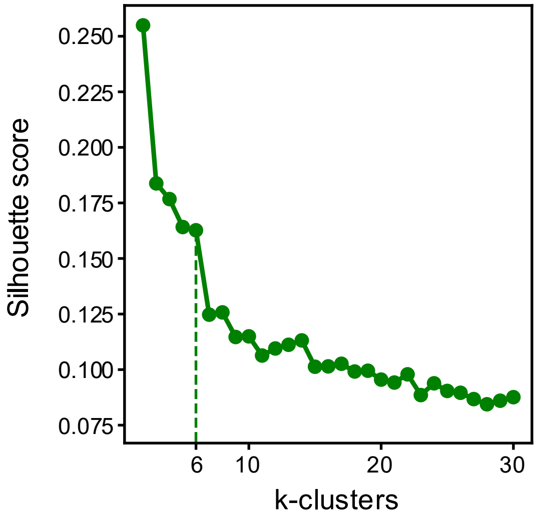


**Supplementary Fig. S11 | Silhouette analysis used to determine the optimal number of K-means clusters.** Silhouette scores for increasing values of *k* are shown, supporting the choice of six clusters for the transcriptomic dataset.

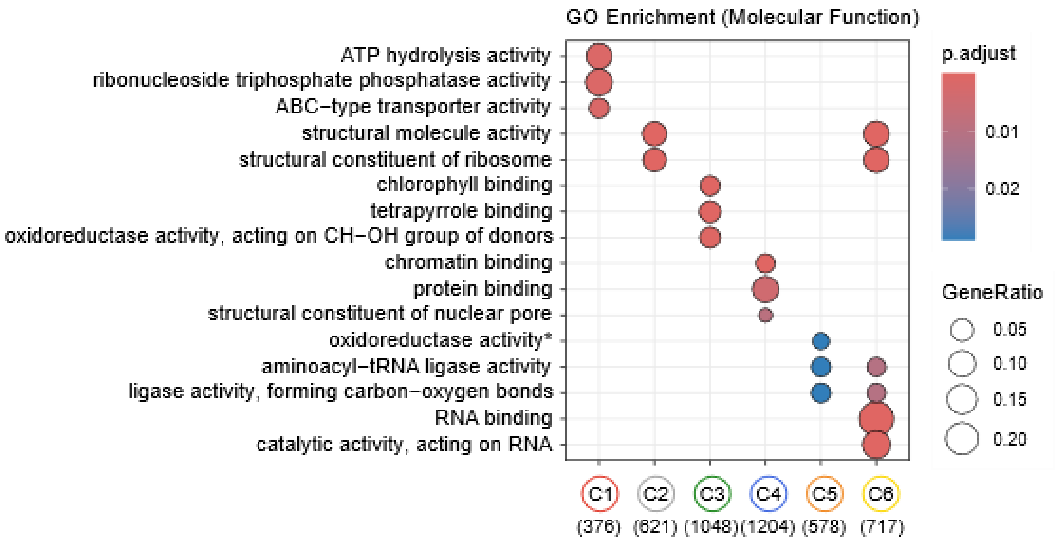


**Supplementary Fig. S12 | Gene Ontology (GO) enrichment per cluster for Molecular Function.** Enriched GO terms (Cellular Component category) are shown for each of the six *K*-means clusters identified in Fig. 4. See Fig. 4 for GO: Biological Process and GO: Cellular Compartment.

### Figure S13A – Cluster 1

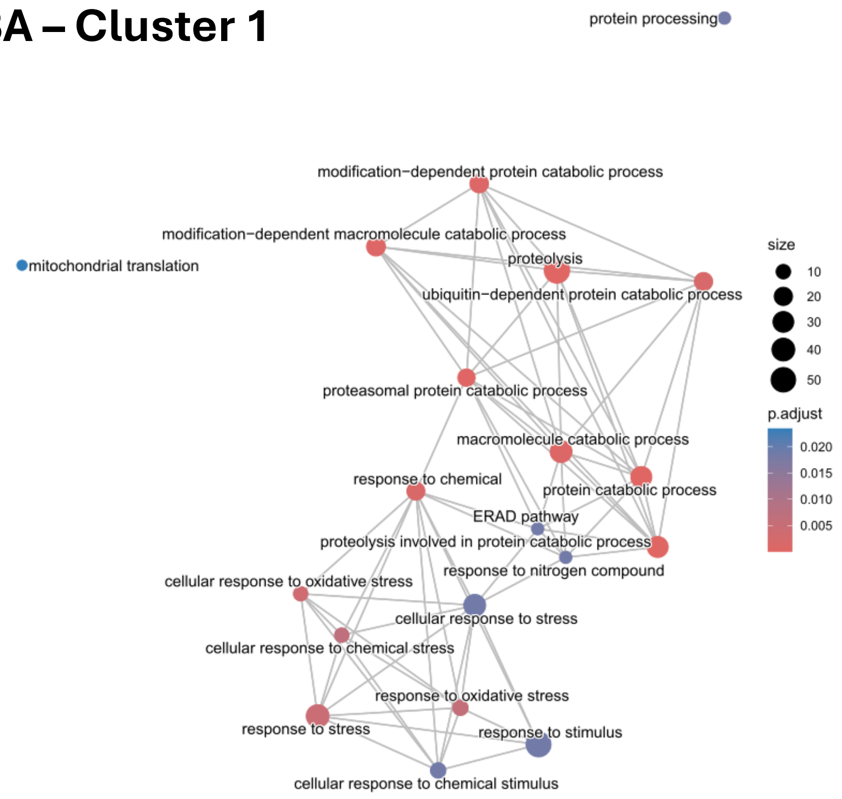


### Figure S13B – Cluster 2

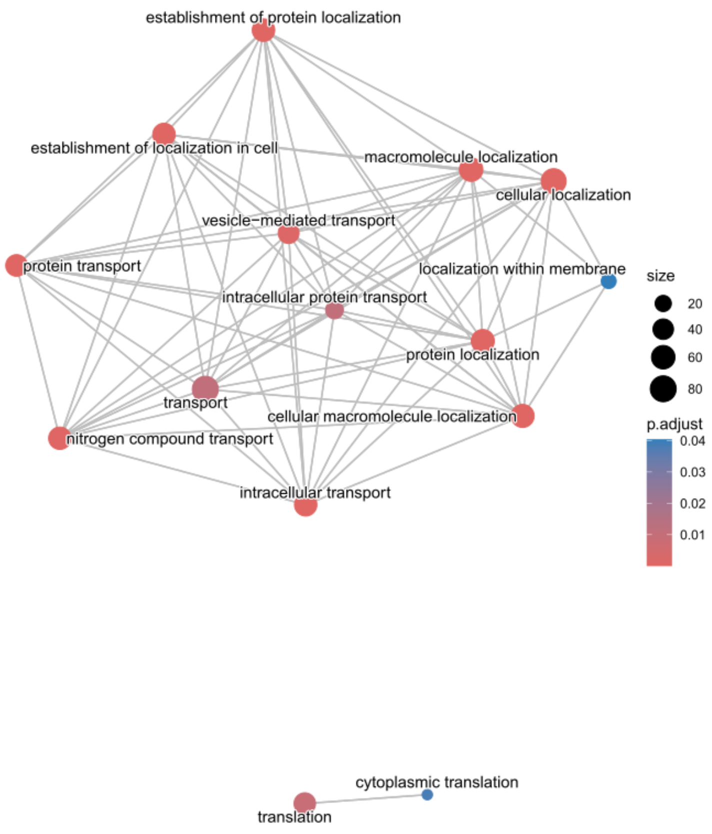

#### Figure S13C – Cluster 3

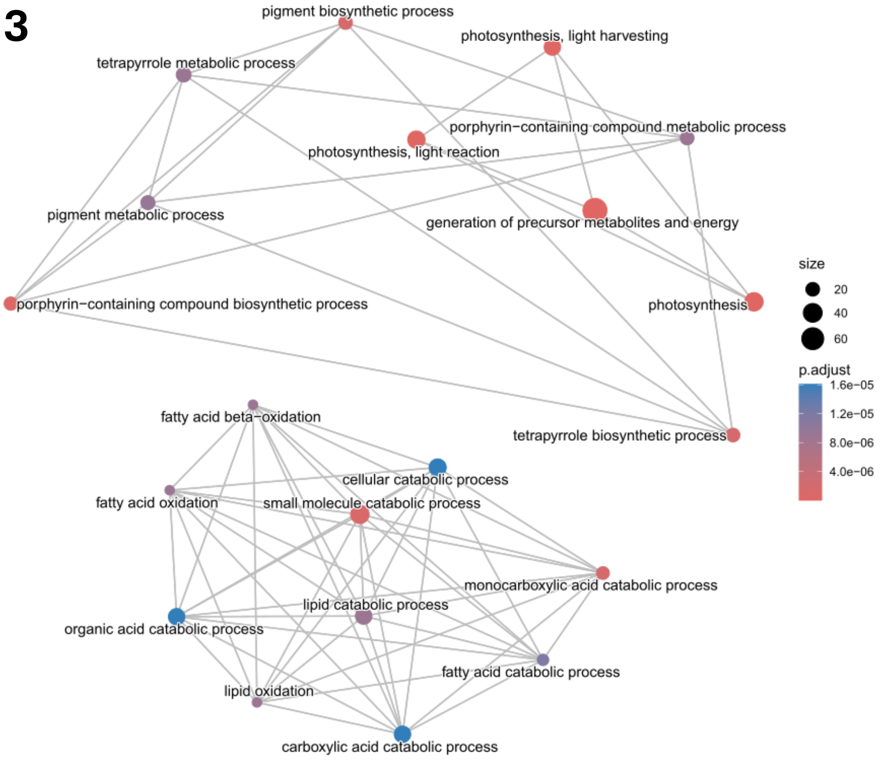


#### Figure S13D – Cluster 5

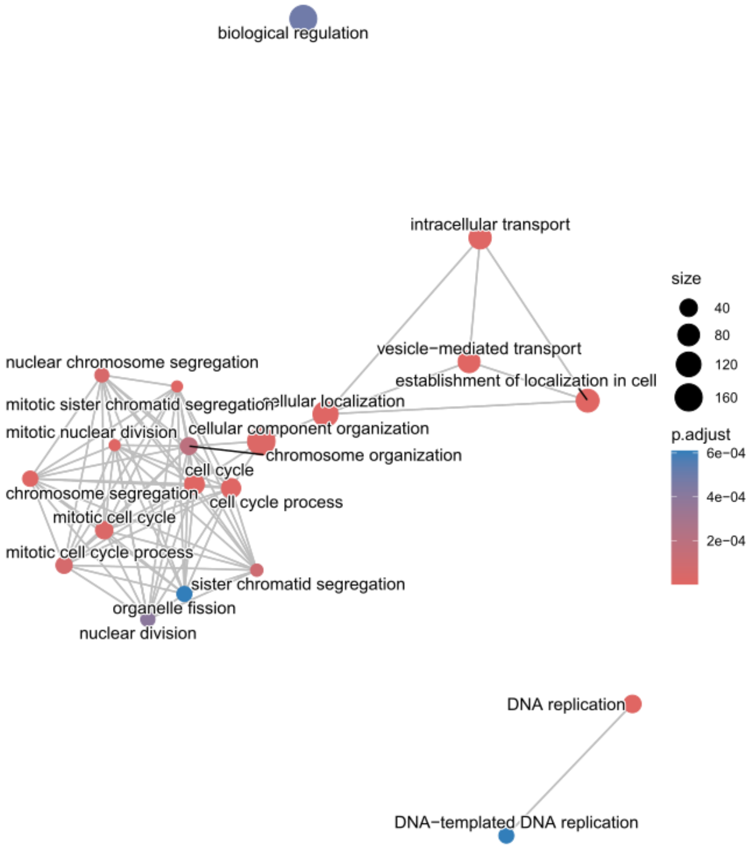

### Figure S13D – Cluster 5

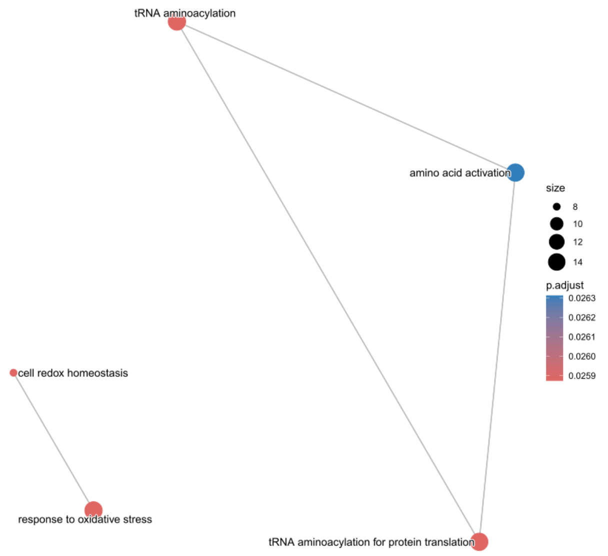


### Figure S13F – Cluster 6

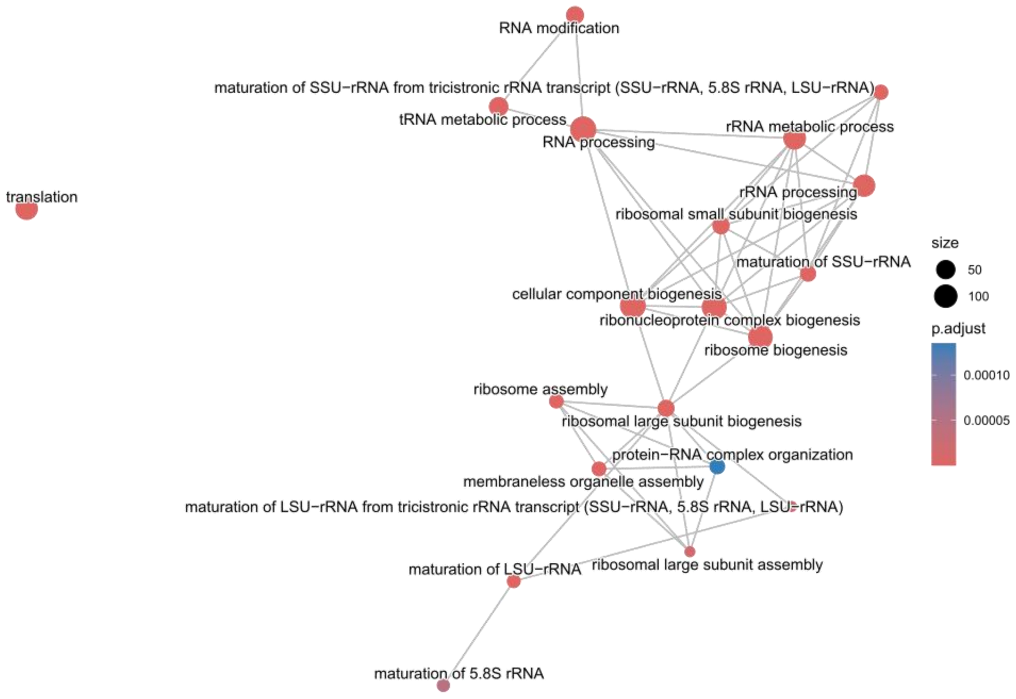


**Fig. S13 | Gene Ontology enrichment networks for each expression cluster.** GO enrichment networks were generated separately for each of the six expression clusters using the *emaplot*() function from the enrichplot R package. Nodes represent significantly enriched GO terms (*P*-value < 0.05)
